# Growth-induced physiological hypoxia correlates with growth deceleration during normal development

**DOI:** 10.1101/2024.06.04.597345

**Authors:** Yifan Zhao, Cyrille Alexandre, Gavin Kelly, Gantas Perez-Mockus, Jean-Paul Vincent

## Abstract

Growth deceleration is a universal feature of growth during development: most organs and tissues slow down their growth rate much before growth termination. Using transcriptomics analysis, we show that during their two-day period of growth deceleration, wing imaginal discs of *Drosophila* undergo a progressive metabolic shift away from oxidative phosphorylation and towards glycolysis. We then develop an ultra-sensitive reporter HIF-1α activity, which reveals that imaginal discs become increasingly hypoxic during development in normoxic conditions, suggesting that limiting oxygen supply could underlie growth deceleration. Growth is energetically expensive and thus expected to contribute, indirectly, to oxygen consumption. Indeed, excess TOR signalling, a key stimulator of growth, triggers hypoxia locally and systemically, highlighting the need to rein in growth when oxygen becomes limiting. This is achieved by a negative feedback loop whereby the classic TOR-inhibitory function of HIF-1α is deployed in response to developmental hypoxia. The absence of Sima/HIF-1α leads to cellular stress, which is alleviated by reduced TOR signalling. Conversely, a small increase in oxygen supply reduces the stress induced by excess TOR activity. We conclude that mild hypoxia is a normal feature of organ development and that Sima/HIF-1α prevents growth-induced oxygen demand from exceeding supply.

## INTRODUCTION

In many instances, growth during development follows an S-shaped curve, with a rapid initial phase followed by an inflexion point and a progressive slowing down. Examples of such growth behaviour have been documented in whole animals, including mammals^1,2,3^, birds^4^, crustaceans^5,6^, insects^7^, and planarian worms^8^, as well as in organs and appendages such as miniature swine livers^9^, human tibias^10^, bird wings^11^ and *Drosophila* eye precursors^12^. Natural populations follow a similar growth pattern, which can be modelled by logistic functions^13^. In this case, growth deceleration can be attributed to limited resources^14^.

However, in a well-fed animal, resources are not expected to be limiting and the basis of growth deceleration is not known. In multicellular organisms, growth is controlled by a wide range of systemic and intrinsic signals. Systemic cues include growth hormones and insulin-like growth factors that coordinate the growth of multiple organs^15^, and modulate final size according to nutrient availability^15,16,17,18^. For example in *Drosophila*, a pulse of the steroid hormone ecdysone has been shown to trigger proliferation arrest at the time of pupariation^19,20^. However, despite the importance of ecdysone signalling for appendage growth, there is no documented evidence that hormonal signalling declines during development. On the contrary, ecdysone titres progressively rise before the pulse that triggers proliferation arrest. Therefore, there is no simple correlation between ecdysone signalling and growth deceleration. Tissue-intrinsic cues from morphogen signalling are also known to contribute to growth control both in *Drosophila* imaginal discs and other tissues ^21,22,23^. However, as with hormonal signalling, morphogen signalling continues to rise, at least in *Drosophila* wing imaginal discs, suggesting that it cannot account for growth deceleration. Finally, theoretical considerations have suggested that mechanical feedback could dampen growth as tissue size increases^24,25,26^.

However, so far, there is no experimental evidence that mechanical constraints govern whole organ growth deceleration. Therefore, despite the universal nature of growth deceleration, its molecular basis remains unknown.

As a step towards uncovering the molecular changes that could underpin growth deceleration, we set out to identify mRNAs that quantitively correlate or anti-correlate with growth rate during development. We chose to do this with wing imaginal discs of *Drosophila*, a well-established model of growth control^21,22,24,27^. Volume measurements confirmed that during the third larval instar (L3), wing disc growth has already begun to decelerate. Bulk RNA-Sequencing (RNA-Seq) showed that mRNAs encoding components of aerobic metabolism become increasingly less abundant with disc age, suggesting that discs may become mildly hypoxic as they enlarge and their growth rate declines. This was confirmed with a highly sensitive reporter of the activity of Sima/HIF-1α, the master regulator of hypoxia response. We then demonstrate that rising Sima/HIF-1α activity causes a reduction in TOR signalling, hence dampening growth. Conversely, excess TOR activity leads to increased hypoxia, and induces cellular stress, presumably by putting impossible demands on oxygen supply. Although one cannot directly demonstrate that limiting oxygen availability accounts for growth deceleration, our results show that the classic hypoxia response mediated by Sima/HIF-1α is deployed during normal development to ensure that oxygen demand imposed by growth does not exceed oxygen supply.

## RESULTS

### Transcriptomic profiling of L3 wing discs suggests a progressive decrease in aerobic metabolism

Wing imaginal discs originate from a group of about 30 cells set aside during embryogenesis^28,29,30^. Over the course of the three larval instars (L1-L3), they undergo 10-11 cell cycles to generate the 30,000-50,000 cells that make up adult wings^31,32^. We measured the volume (a proxy for tissue mass) of precisely timed wing discs, from the onset of L3, at 72 hours after egg laying (AEL) (Fig. 1a & 1b). Wing disc volumes, expressed as a ratio to starting volume, were plotted over time (Fig. 1c) and the relative growth rate was determined as its first derivative, normalised to the instantaneous volume. This analysis confirmed that the relative growth rate progressively decelerates, declining from 9% per hour at the beginning of L3 to 2% per hour at the end of L3, before pupariation (Fig. 1d).

**Fig. 1:**
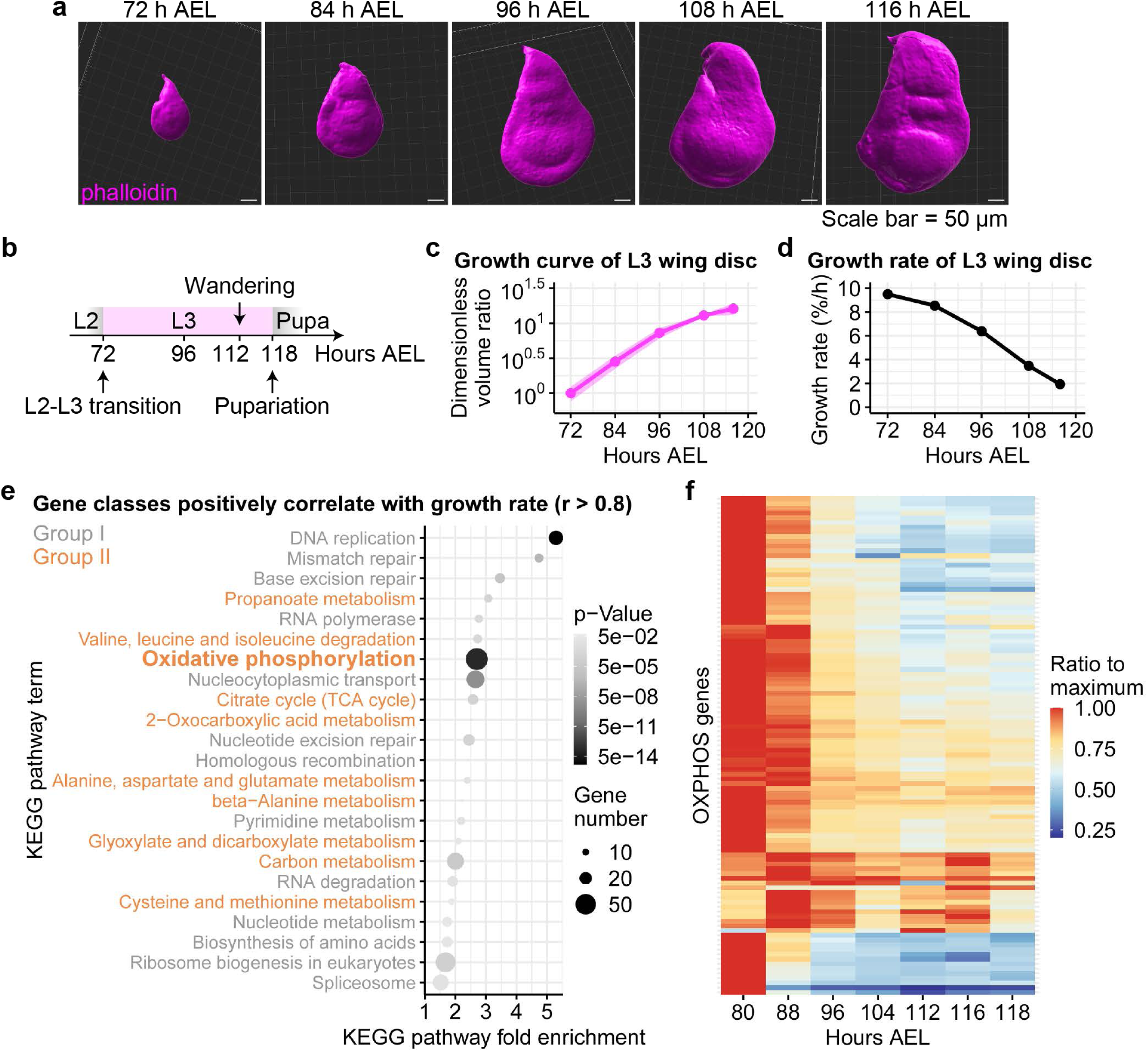
Transcriptomic analysis of L3 wing discs suggests progressive decrease in aerobic metabolism. **a.** Representative volumetric reconstruction of phalloidin-stained L3 wing imaginal discs. Confocal images were taken and reconstructed under identical conditions (n = 6 wing discs for each time point, except for data at 84 hours AEL where n = 5). **b.** Schematic timeline of development during the third instar (72 to 117 hours AEL). **c.** Plot of imaginal disc volumes during L3. Dimensionless volumes (normalised to the average initial volume are shown (n = 6 wing discs for each time point, except for data at 84 hours AEL where n = 5, error ribbon represents standard deviations). **d.** Plot of disc growth rate, calculated as the first derivative of the growth curve (*dV/dt*) divided by the volume (*V*) at any given time point. **e.** KEGG pathway analysis of genes with expression profiles that positively correlate with the growth rate (r > 0.8). Central dogma genes (Group I) and aerobic respiration (Group II) are downregulated overtime, with DNA replication and OXPHOS being the most significantly enriched. **f.** Transcriptional profile of OXPHOS genes expressed during L3. For each gene, the expression level was normalised to the maximum value, with the normalised expression color-coded according to the lookup table on the right (n = 3 biological replicates for each time point, except for data at 80 hours AEL where n = 2).

We then sought to identify mRNAs that correlate or anti-correlate with the instantaneous growth rate. Wing discs grow relatively homogenously^33^, alleviating the need for spatially resolved analysis. Bulk RNA-Seq was performed on precisely timed L3 wing discs at seven developmental time points, ranging from 80 to 118 hours AEL. For each expressed gene, a time-course expression profile was generated and the Pearson correlation coefficient (r) with growth rate was obtained. Genes with an absolute r value exceeding 0.8 were considered of interest. They were analysed using the Kyoto’s Encyclopedia of Genes and Genomes (KEGG) database resource^34^ to reveal the molecular functions of mRNAs that correlate with growth rate, i.e. that decrease with increased tissue size (Fig. 1e). Two groups of genes stood out. Group I encompasses genes associated with DNA, RNA and protein synthesis, activities that are required for biomass accumulation. The presence of these genes among our hits is expected from growth deceleration; they will therefore not be considered further. Group II comprises genes involved in oxidative phosphorylation (OXPHOS), tricarboxylic acid (TCA) cycle and acetyl coenzyme A (acetyl-CoA) biosynthesis (Fig. 1f, Extended Data Fig. 1a-c), suggesting a progressive decrease in aerobic metabolism as wing discs grow towards their final size. In addition, our analysis revealed a significant increase in the expression of many genes encoding glycolytic enzymes, including *phosphofructokinase* (*Pfk*), *aldolase 1* (*Ald1*), *triose-phosphate isomerase* (*Tpi*), *glyceraldehyde-3-phosphate dehydrogenase 2* (*Gapdh2*) and *phosphoglycerate kinase* (*Pgk*) (Extended Data Fig. 1d). Of note, mRNA encoding *Glucose transporter 1* (*Glut1*) was found to increase 14-fold during L3 (Extended Data Fig. 1d), suggesting an increase in anaerobic metabolism. A decrease in OXPHOS combined with an increase in anaerobic metabolism is reminiscent of the response to hypoxia^35,36,37,38,39,40,41^. Accordingly, we detected a small rise in the activity of an enhancer trap^42,43^ inserted in the gene encoding *lactate dehydrogenase* (*Ldh*), a classic hypoxia-induced gene^44^ (Extended Data Fig. 1e & 1e’). We next set out to assess the level of hypoxia in wing discs with dedicated sensors.

### Developing L3 wing discs become increasingly hypoxic in environmental normoxia

A previously described transgenic oxygen sensor termed nlsTimer takes advantage of a fluorophore that matures to two distinct isoforms, with a green variant being favoured at low oxygen tension and a red one at high oxygen tension^45^ (Extended Data Fig. 2a). Thus, the red-to-green ratio after a pulse of expression gives an indication of oxygen level. We triggered expression of this sensor in whole larvae at 72 and 84 hours AEL and performed ratiometric fluorescence analysis 32 hours later (the time needed for maturation of the fluorophores) (Extended Data Fig. 2b). The results suggest that whole larva oxygen levels could indeed decrease as development proceeds (Extended Data Fig. 2c, 2c’ & 2c’’).

Unfortunately, we were not able to estimate cellular oxygen specifically in imaginal discs because the sensor had an adverse effect on imaginal disc growth (Extended Data Fig. 2d). Therefore, the activity of nlsTimer is suggestive of mild hypoxia in late L3 but one cannot infer whether this is reflected in reduced oxygen in imaginal discs.

Another means of estimating oxygen level is by leveraging the mechanism whereby oxygen tension modulates the activity of Hypoxia Inducible Factor-1α (HIF-1α)^46^, a transcription factor known as Similar (Sima) in *Drosophila*^47^. Under normoxia, *Drosophila* HIF prolyl hydroxylase (Hph, AKA Fatiga; PHD in mammals) hydrolyses Sima/HIF-1α at specific proline residues within the so-called oxygen-dependent degradation domain (ODD)^48,49,50,51,52^, thus earmarking it for ubiquitylation by the von Hippel Lindau (VHL) protein and degradation by the proteasome^53,54,55^. Under hypoxia, Sima/HIF-1α hydroxylation and hence ubiquitylation is reduced^50^, leading to its accumulation in the nucleus and activation of target genes^56^. ODD confers oxygen-modulated degradation to heterologous proteins (e.g. GFP), and this is the basis of another oxygen biosensor, GFP-ODD^57^. Although this reporter is detectably activated in larvae cultured in 5% environmental oxygen^58^, we found that its activity, normalised to a reference mRFP-producing transgene, did not differ noticeably between early and late L3 discs (Extended Data Fig. 2e & 2e’), perhaps because of limited dynamic range. Indeed, in *Drosophila* embryos, the average GFP-ODD-to-mRFP ratio only showed a 2-fold increase from 60% to 5% environmental oxygen^57^.

While sensitive detection of oxygen tension is difficult to achieve *in vivo*, Sima/HIF-1α activity can, with caveats, be used as a proxy for oxygen level within cells. Indeed, several reporters comprising consensus Sima/HIF-1α binding sites upstream of a minimal promoter and a marker gene have been generated^52,59,60,61,62,63^. For example, a Lac-Z reporter based on a murine 233-bp hypoxia-responsive enhancer comprising two consensus hypoxia response elements (HREs, 5’-(A/G)CGTG-3’) and a cAMP response element was found to be activated at 5% environmental oxygen in transgenic *Drosophila* embryos^52^. However, this reporter showed no activity in milder hypoxic condition^64^, suggesting that improvements in sensitivity are needed. We reasoned that this could be achieved with a native hypoxia-responsive enhancer and by taking advantage of recent improvements in reporter gene design in *Drosophila*^20^. As a hypoxia-responsive enhancer, we used an 81-bp fragment from the *Drosophila hph* gene comprising two putative HREs - HRE2 and HRE3 - which were characterised in S2 cells by Wappner and colleagues^65^. Five tandem copies of this fragment were inserted upstream of a minimal heat shock promoter driving the production of mRNA encoding nuclear targeted NeonGreen tetramers (NLS4xNG). Viral untranslated sequences (syn21 at the 5’ and p10 at the 3’) were included to boost expression^66^ (Fig. 2a).

**Fig. 2:**
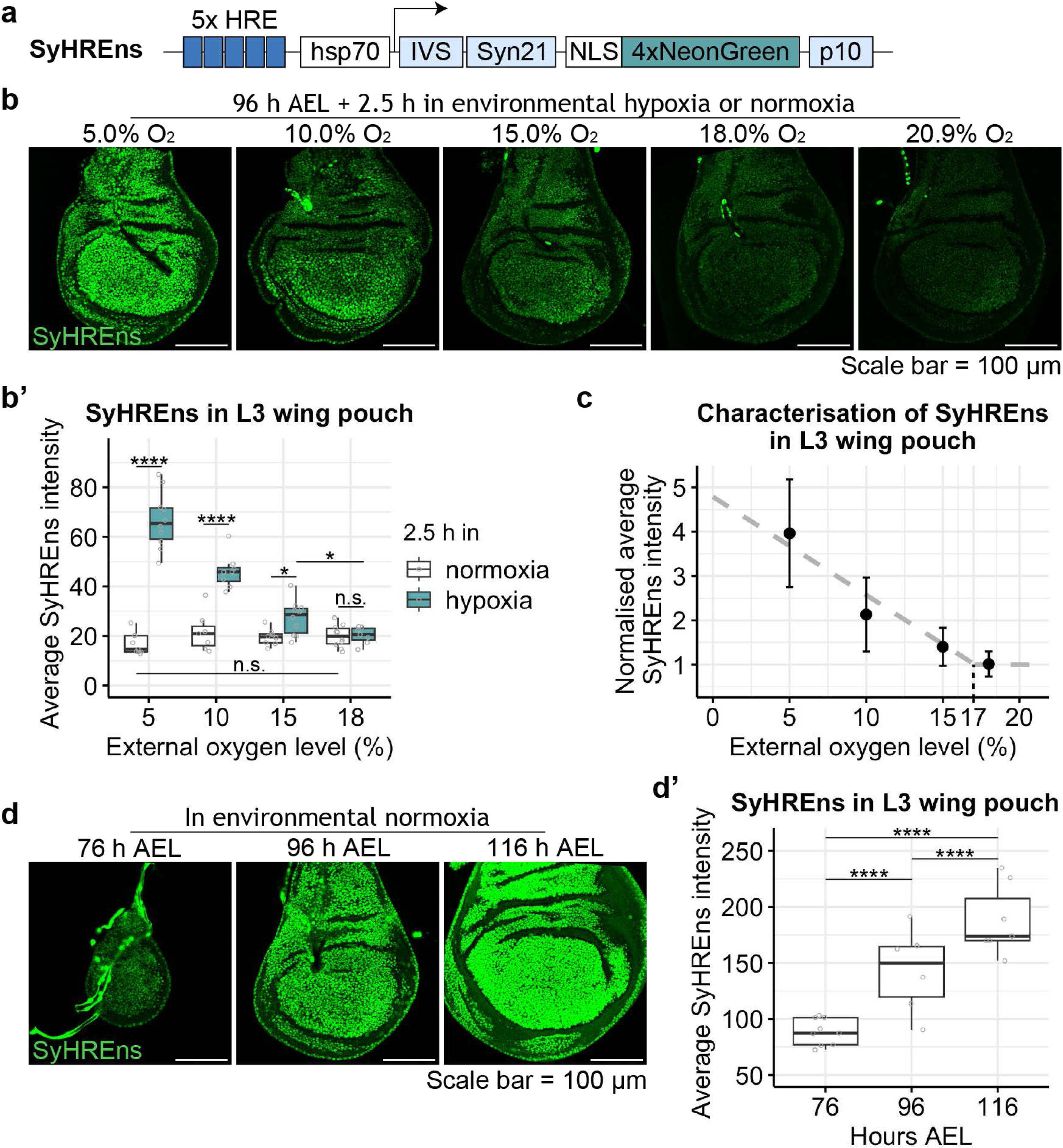
Developing L3 wing discs become increasingly hypoxic in environmental normoxia. **a**. Schematic representation of the hypoxia reporter, which we have named SyHREns. Note that the presence of two SyHREns transgenes led to mild developmental delay (one hour over the approximately five days to pupariation) but had no other detectable effect on development. **b-b’**. SyRHEns fluorescence intensity in imaginal discs dissected from larvae cultured in different environmental oxygen tension for 2.5 h. Representative images are shown in **b** and quantification in **b’** (n ≥ 7 wing discs for each time point). After ensuring normality of data with a Shapiro-Wilk test, a two-sided unpaired t-test was used for statistical significance (* for p < 0.05, **** for p < 10^-4, n.s. for not significant). For all the box plots in this and subsequent figures, the centre line denotes the median (50th percentile); the lower quartile (Q1) denotes the 25th percentile; the upper quartile (Q3) denotes the 75th percentile; the interquartile range (IQR) is defined as *Q*3 − *Q*1; the outliers are defined as data points that lie below *Q*1 − 1.5 × *IQR* or above *Q*3 + 1.5 × *IQR*. **c**. Linear regression of the SyHREns intensity versus external oxygen (error bars represent standard deviations). **d-d’**. SyHREns activity progressively rises in L3 wing discs under environmental normoxia. Representative images are shown in **d** and quantification in **d’** (n ≥ 7 wing discs for each time point, except for data at 96 hours AEL where n = 6). After ensuring the normality of the data, a two-sided unpaired t-test was performed to assess statistical significance (**** for p < 10^-4).

To assess the sensitivity and dynamic range of the resulting construct, called SyHREns (Synthetic HRE nuclear sensor) for the remainder of this paper, transgenic larvae from mid-L3 (96 hours AEL) were cultured for 2.5 hours in different levels of oxygen, and the imaginal discs were then dissected out and imaged by fluorescence microscopy (Fig. 2b). The result showed that hypoxia activates the reporter in a severity-dependent manner (Fig. 2b’). A plot of reporter activity at various oxygen levels (normalised to that in normoxia) suggests that a relatively mild departure from normoxia (17% oxygen) can be detected and that the response is relatively linear, with a 4-fold signal increase at 5% oxygen (Fig. 2c). These observations suggest that SyHREns is highly sensitive with an excellent dynamic range and therefore suitable to infer any change in oxygen level during normal development. From early to late L3, a gradual increase in SyHREns fluorescence was detected both in wing (Fig. 2d & 2d’) and eye discs (Extended Data Fig. 2f & 2f’), indicating a progressive increase in Sima/HIF-1α activity. The inference that this is due to hypoxia is reinforced by the behaviour of nlsTimer, a direct reporter of oxygen level (Extended Data Fig. 2c, 2c’ & 2c’’). We conclude that wing imaginal discs become increasingly hypoxic as they grow during L3.

### Sima/HIF-1α dampens TOR signalling through activation of Scylla/REDD1

In *Drosophila*, as in mammals, excess Sima/HIF-1α causes a reduction in cell size and number^58,67,68,69^. We confirmed that overexpression of *sima/HIF-1α* in the posterior (P) compartment of imaginal discs leads to marked tissue size reduction (Fig. 3a). This was accompanied by a large increase in SyHREns activity, as expected. A milder but similar effect was seen upon *hph* knockdown, which is expected to cause accumulation of Sima/HIF-1α.

**Fig. 3:**
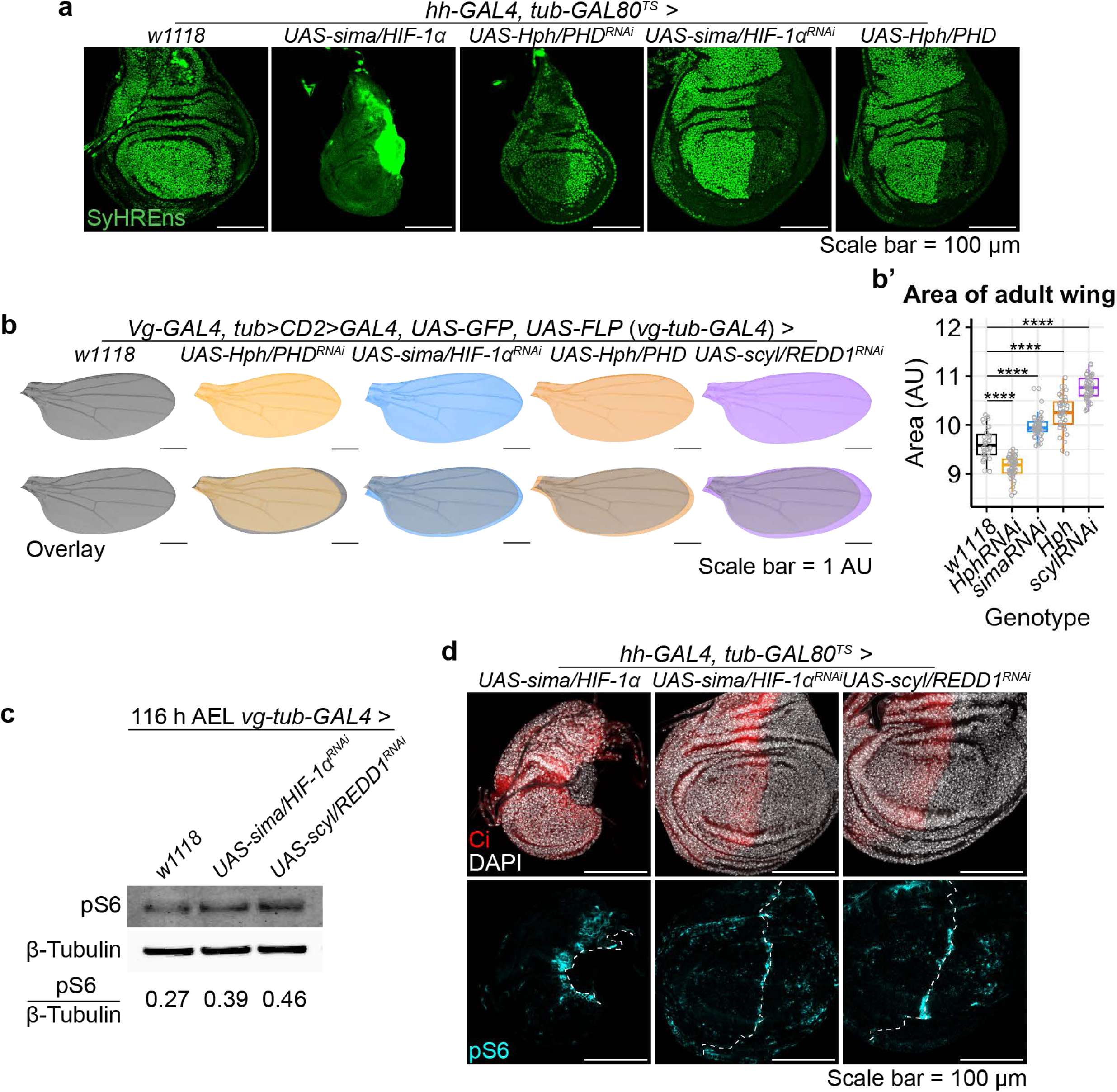
Sima/HIF-1α dampens growth by suppressing TOR activity through transcriptionally activation of *scyl/REDD1*. **a**. Perturbation of *sima/HIF-1α* or *Hph/PHD* in the posterior compartment (with *hedgehog* (*hh*)*-GAL4*) affects SyHREns activity. **b-b’**. Whole disc perturbations of *sima/HIF-1α*, *Hph/PHD* or *scyl/REDD1* (with *vg-tub-GAL4*) affects adult wing size. Representative images are shown in **b** and quantification in **b’** (n ≥ 42 adult wings for each genotype, except for *w1118* control wings where n = 32). After ensuring the normality of the data with a Shapiro-Wilk test, a two-sided unpaired t-test was used to assess statistical significance (**** for p < 10^-4). **c**. Western blot analysis shows a mild increase in pS6 upon whole disc knockdown of *sima/HIF-1α* or *scyl/REDD1* (*vg-tub-GAL4 > sima/HIF-1α^RNAi^* or *scyl/REDD1^RNAi^*). The pS6 signal, normalised with anti-β-Tubulin is indicated as a ratio under each lane. **d**. Effect of modulating *sima/HIF-1α* or *scyl/REDD1* activity on pS6 immunoreactivity. Overexpression of *sima/HIF-1α* (*hh-GAL4 > sima/HIF-1α*) dampens the pS6 signal while *sima/HIF-1α* or *scyl/REDD1* knockdown (*hh-GAL4 > sima/HIF-1α^RNAi^* or *scyl/REDD1^RNAi^*) enhances it. The domain of *hh-GAL4* expression is marked by the absence of Ci (red). High pS6 immunoreactivity at the A-P boundary (particularly visible in the left-handed panel) is also noticeable in wild type, an observation that we do not rationalise.

Genetic manipulations that <u>reduce</u> Sima/HIF-1α activity had the expected inhibitory effect on SyHREns but did not noticeably affect tissue size in wing imaginal discs. To further assess whether reducing Sima/HIF-1α activity promotes excess growth, we examined adult wings arising from imaginal discs that experienced *sima/HIF-1α* knockdown or *hph* overexpression during L3. A small but statistically significant increase in surface area was seen in both cases (Fig. 3b & 3b’). We conclude that, as expected from previous evidence, Sima/HIF-1α dampens tissue growth while loss of activity only leads to mild overgrowth.

In mammalian cells, hypoxia-induced HIF-1α activity reduces growth by suppressing the activity of the mammalian Target of Rapamycin (mTOR)^70,71,72,73,74^, a major regulator of anabolic metabolism^75^. It is of interest therefore that TOR signalling activity, as assayed by phosphorylated ribosomal protein S6 (pS6) immunoreactivity, progressively decreases during the growth of L3 wing imaginal discs^76,77^, an observation that we confirmed by Western blot analysis (Extended Data Fig. 3a & 3a’). We also detected a progressive decline in O-propargyl-puromycin (OPP) incorporation in L3 wing discs (Extended Data Fig. 3b & 3b’), an indication of decreasing rate f protein synthesis. Therefore, during L3 disc development, TOR signalling and protein synthesis decrease, while Sima/HIF-1α activity rises. To test whether there is a causal relationship between these two trends, we asked whether modifying Sima/HIF-1α activity had any effect on pS6. Western blot analysis revealed that, upon whole disc *sima/HIF-1α* knockdown, pS6 immunoreactivity (normalised to β-Tubulin) is higher than normal at the end of larval development (Fig. 3c & Extended Data Fig. 3c).

Similarly, compartment-specific knockdown also led to higher pS6 immunoreactivity than in the control A compartment (Fig. 3d, middle column). And conversely, *sima/HIF-1α* overexpression suppressed S6 phosphorylation (Fig. 3d, left-handed column). Taken together, these results confirm the expectation that Sima/HIF-1α downregulates TOR signalling activity.

Studies in *Drosophila* and in mammalian cells have shown that regulation of TOR signalling by hypoxia requires REDD1, known as Scylla (Scyl) in *Drosophila*^70,78^. We therefore tested whether, in environmental normoxia, Scyl/REDD1 mediates the inhibitory effect of Sima/HIF-1α on TOR signalling. RNAi against *scyl/REDD1* in wing imaginal discs leads to a 12% enlargement of adult wings (Fig. 3b & 3b’) as well as an increase pS6 immunoreactivity, on par with that seen upon *sima/HIF-1α* knockdown (Fig. 3c, 3d & Extended Data Fig. 3c). These observations confirm that Scyl/REDD1 normally restrains growth by dampening TOR activity. Our RNA-Seq dataset shows a 5-fold increase in *scyl/REDD1* mRNAs during L3 (Extended Data Fig. 3d), paralleling the rise in HIF-1α activity. This raises the possibility that Sima/HIF-1α activates *scyl/REDD1* transcription. Indeed, *sima/HIF-1α* overexpression caused a marked increase in *scyl/REDD1* RNA levels, as assayed *in situ* by hybridisation chain reaction (HCR)^79^ (Extended Data Fig. 3e, middle column). The converse, loss of *scyl/REDD1* transcripts in response to *sima/HIF-1α* knockdown, is also true although less obvious because of the relatively low basal expression of *scyl/REDD1* in the prospective wing area at L3 (Extended Data Fig. 3e). Overall, the above findings suggests that during normal disc growth, rising Sima/HIF-1α activates Scyl/REDD1, which in turn dampens growth by down-regulating TOR activity.

### Excess TOR activity leads to autonomous and non-autonomous hypoxia

Rapid growth requires ATP, normally released by the breakdown of nutrients in the presence of oxygen. It is expected therefore that, unless ATP production enlists inefficient anaerobic catabolism, the demand of growth for ATP would, indirectly, put pressure on oxygen supply.

We examined this possibility by asking whether excess TOR activity affects Sima/HIF-1α activity. TOR activity was boosted specifically in the P compartment by overexpressing *Ras homolog enriched in brain* (*Rheb*) with *UAS-Rheb* and *hh-GAL4*. This led to a marked autonomous increase in SyHREns fluorescence intensity (Fig. 4a & 4a’), consistent with the notion that excess TOR signalling triggers hypoxia. To exclude the trivial explanation that increased SyHREns fluorescence arises from TOR-induced increased translation of the reporter, we assessed the effect of *Rheb* overexpression on the fluorescence from a *ubiquitin-NLS4xNeonGreen* (*ubi-NLS4xNG)* transgene, which is not expected to respond directly to oxygen levels. A mild increase of fluorescence was detected in the *Rheb* overexpression domain (Extended Data Fig. 4a & 4a’), suggesting that increased translation does contribute to the Rheb-induced SyHREns signal, albeit in a relatively modest fashion. It appears therefore that the bulk of the SyHREns signal is indeed due to hypoxia. As an independent assay for the effect of excess TOR activity on oxygen levels, we returned to the GFP-ODD reporter, which has poor sensitivity but has the benefit of estimating oxygen levels from a fluorescence ratio and is therefore independent of protein synthesis levels. The results confirmed that *Rheb* overexpression leads to cell autonomous hypoxia (Extended Data Fig. 4b, 4b’ & 4b’’), suggesting that TOR activity puts pressure on oxygen supply.

**Fig. 4:**
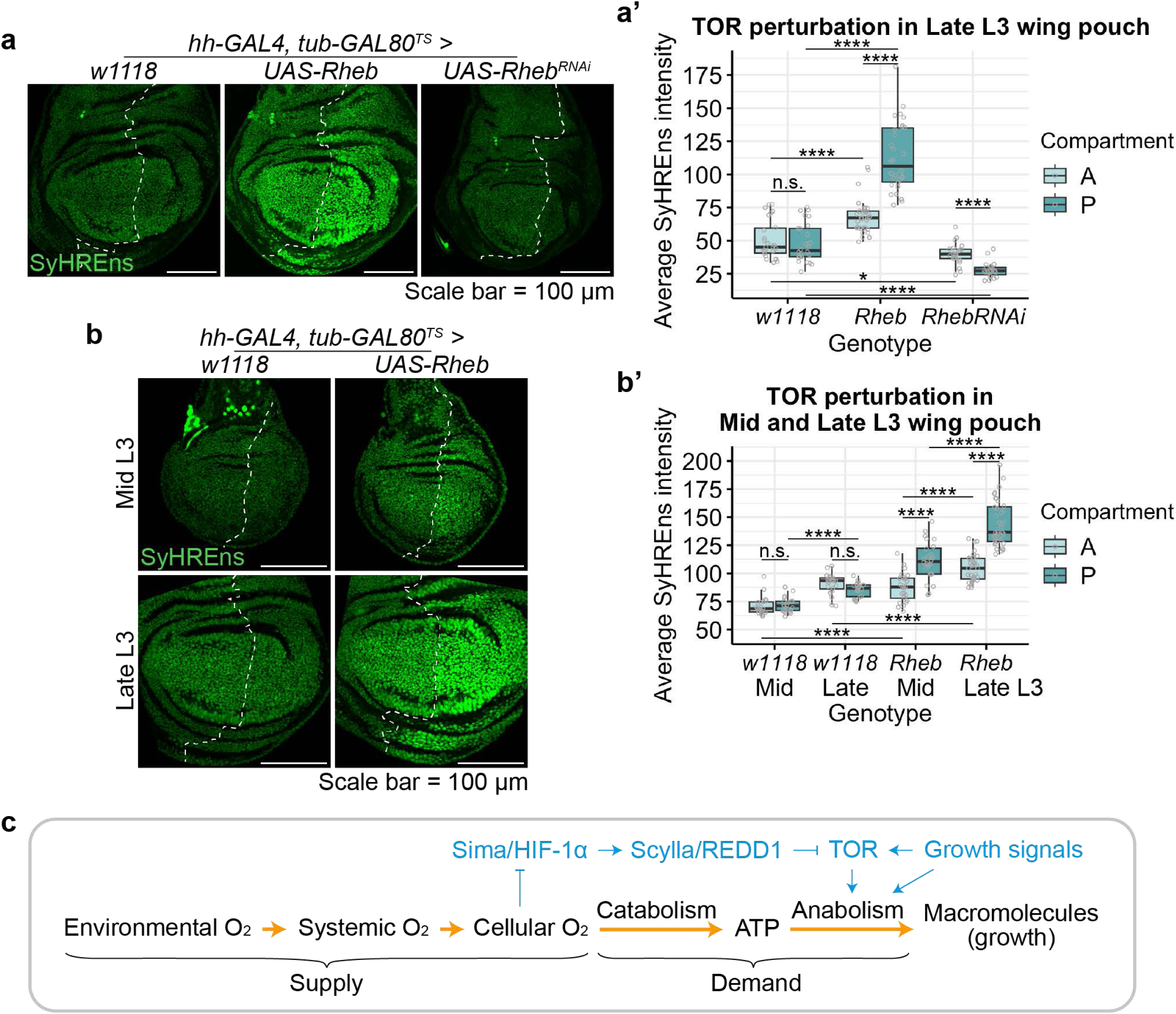
Excess TOR activity leads to autonomous and non-autonomous hypoxia. **a-a’**. *Rheb* overexpression in the P compartment boosts SyHREns activity most prominently in the P compartment but also non-autonomously in the A compartment (as seen by comparing fluorescence intensity in the A compartments of experimental (middle panel) and control discs (left-handed panel). *Rheb* knockdown has the opposite effect on SyHREns activity, also autonomously and non-autonomously. Representative images are shown in **a** (all discs imaged under identical conditions) and quantification in **a’** (n ≥ 13 wing discs for each genotype, except for data of *hh-GAL4, tub-GAL80^TS^* > *UAS-Rheb^RNAi^* where n = 11). After testing for normality with a Shapiro-Wilk test, a two-sided unpaired Wilcoxon signed-rank test was used to assess statistical significance (* for p < 0.05, **** for p < 10^-4, n.s. for not significant). **b-b’**. The autonomous and non-autonomous effect of *Rheb* overexpression on SyHREns activity is more marked in late L3 than in mid-L3 discs. Representative images are shown in **b** and quantification in **b’** (n ≥ 10 wing discs for each condition). Depending on the normality of the data, a two-sided unpaired t-test or a two-sided unpaired Wilcoxon signed-rank test was used to assess statistical significance (**** for p < 10^-4, n.s. for not significant). **c**. The role of oxygen and Sima/HIF-1α in mediating negative feedback of growth onto itself. Environmental oxygen is transported into the organism before being distributed to cells. TOR signalling reduces cellular oxygen by promoting anabolism, and dwindling oxygen, in turn, suppresses TOR signalling via the Sima/HIF-1α - Scyl/REDD1 axis (blue).

As shown in Fig. 2d, the hypoxic state of imaginal discs rises progressively with age, suggesting that oxygen could be more limiting in old (large) than in young (small) discs. To investigate this possibility, we compared the effect of excess TOR activity on SyHREns activity in mid and late L3 discs. But first, we examined the effect of Rheb-induced translation on the control *ubi-NLS4xNG* transgene at mid and late L3. A similarly mild increase was seen at both stages, and only in a cell autonomous fashion (Extended Data Fig. 4c & 4c’). In contrast, Rheb had a much more marked effect on SyHREns activity at late L3 than at mid-L3 (Fig. 4b & 4b’). We conclude that the pressure on oxygen supply rises with tissue age/size. This may explain why oxygen supplementation is required for the growth of cultured imaginal discs^80^.

The cell autonomous effect of excess TOR signalling on SyHREns activity (as shown in *hh- GAL4*, *UAS-Rheb*) is particularly striking. However, a milder effect can also be detected in the genetically unaffected A compartment (compare the fluorescence to that in the A compartment of control discs that do not overexpress *Rheb*) (Fig. 4a, 4a’, 4b & 4b’). No such non-autonomous effect can be seen with ubi-NLS4xNG, suggesting that overactivation of TOR in the P compartment has no noticeable effect on translation rate in the A compartment (Extended Data Fig. 5c & 5c’). We suggest therefore that excess TOR activity in the P compartment has an adverse effect on oxygen availability in the A compartment, possibly because excess consumption in the P compartment draws oxygen away from the A compartment. We note that the non-autonomous rise in SyHREns activity in the A compartment is relatively uniform, suggesting that the overgrowing tissues draws oxygen broadly, perhaps at the systemic level.

### Excess TOR activity leads to cellular stress in the absence of Sima/HIF-1α

Our results so far suggest that oxygen becomes limiting during growth and that, in response, Sima/HIF-1α reduces oxygen demand by dampening growth. One can wonder why such a mechanism needs to be deployed since, irrespective *of* Sima/HIF-1α activity, reduced oxygen is bound to slow down aerobic metabolism and hence the production of ATP needed for growth. To address this question, we studied the effect of TOR overactivation in the presence or absence of Sima/HIF-1α. Overexpression of *Rheb*, e.g. in the P compartment, led to a cell autonomous increase in tissue size, as expected, but also cellular stress, as indicated by the activity of TRE-NLS4xNG, a reporter of c-Jun N-terminal kinase (JNK) signalling activity (Extended Data Fig. 5a, Fig. 5a & 5a’). This effect was further exacerbated by *sima/HIF-1α* knockdown (Fig. 5a & 5a’). *Sima/HIF-1α* knockdown alone was sufficient to trigger a modest induction of JNK signalling activity, which could be fully rescued by inhibiting TOR activity (Fig. 5b & 5b’). These observations suggest that, if growth makes demands on ATP that cause catabolic pathways to draw more oxygen than is available, tissue stress ensues. The role of Sima/HIF-1α in reining in TOR signalling was further investigated in *sima/HIF-1α* mutant animals. A mutant allele, *sima/HIF-1α*^*KG07607*^ had previously been generated^67^. In addition, we generated a conditional allele, *sima/HIF-1α^>E2-6>^*, which can be converted by FLP recombinase to *sima/HIF-1α^ΔE2-6^,* a null mutant. Both *sima/HIF-1α^KG07607^* and *sima/HIF-1α^ΔE2-6^* were viable over a deficiency albeit with a phenotype characterised by wing crumpling, loss of wing hairs and altered hair shape (Extended Data Fig. 5b, 5c & 5c’). Approximately 75% of homozygous *sima/HIF-1α^ΔE2-6^* adults had at least one crumpled wing. Reduced hair density (a proxy for cell density^81^) was observed at the surface of non- crumpled wing (Extended Data Fig. 5c & 5c’) suggesting an increase in cell size, possibly as a result of excess TOR signalling. The crumpling phenotype was partially rescued by addition of rapamycin in the larval food (Fig. 5d), confirming that it is caused by excess TOR activity, possibly by drawing more oxygen than is available. To test the latter suggestion, we sought to test if the crumpling phenotype could be alleviated by increasing oxygen tension. We found 25% oxygen tension to be toxic, even in wild type animals, presumably because of oxidative stress (Extended Data Fig. 5d). However, 22.5% oxygen was well tolerated and this mild departure from normoxia was sufficient to rescue Rheb-induced stress (Fig. 5e & 5e’). Conversely, mild environmental hypoxia enhanced Rheb-induced stress (Extended Data Fig. 5e & 5e’). These findings suggest that the TOR-induced cellular stress is due to a mismatch between oxygen demand and supply.

**Fig. 5:**
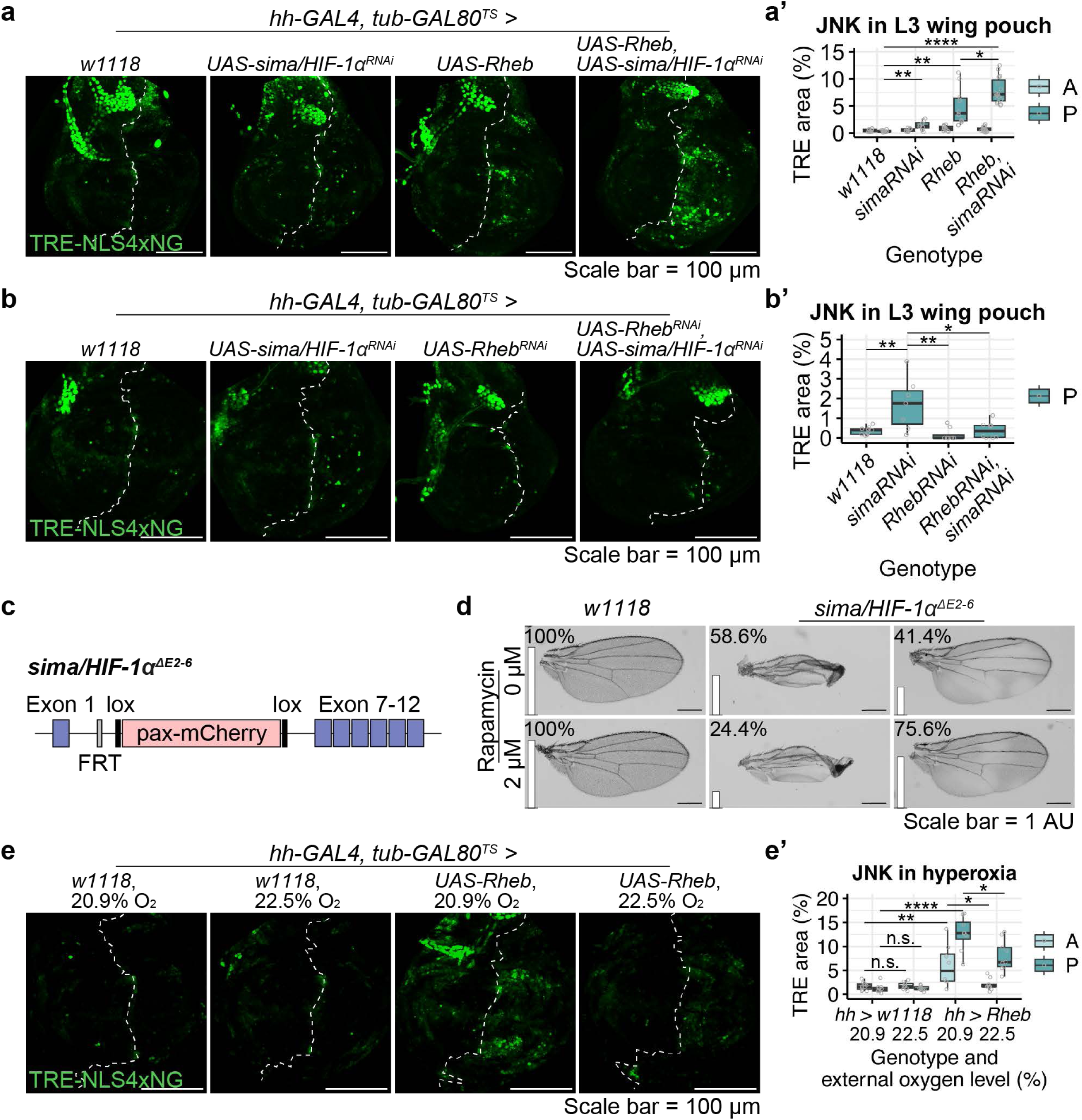
Sima/HIF-1α reins in cellular stress caused by excess TOR activity. **a-a’**. Sima/HIF-1α knockdown or TOR overactivation causes cellular stress (detected by TRE-NLS4xNG). Representative images are shown in **a** and quantification in **a’** (n ≥ 9 wing discs for each genotype, except for data of *hh-GAL4, tub-GAL80^TS^* > *UAS-sima^RNAi^*where n = 8). After ensuring the normality of the data with a Shapiro-Wilk test, a two-sided unpaired t-test was used to assess statistical significance (* for p < 0.05, ** for p < 0.01, **** for p < 10^-4). **b-b’**. Reducing TOR signalling (*Rheb* knockdown) alleviates cellular stress (TRE-NLS4xNG) caused by Sima/HIF-1α knockdown. Representative images are shown in **b** and quantification in **b’** (n ≥ 8 wing discs for each genotype, except for data of *hh-GAL4, tub-GAL80^TS^* > *UAS-sima^RNAi^*where n = 7). Depending on the normality of the data, a two-sided unpaired t-test or a two-sided unpaired Wilcoxon signed-rank test was used to assess statistical significance (* for p < 0.05, ** for p < 0.01). **c**. Schematic representation of the *sima/HIF-1α* KO allele, *sima/HIF-1α^ΔE2-6^*. **d**. Reducing TOR signalling (Rapamycin feeding) partially rescues the wing phenotype of *sima/HIF-1α* KO flies. Representative images and quantification are shown (n ≥ 150 control, w*1118,* wings and n ≥ 78 *sima/HIF-1α^ΔE2-6^* homozygous wings). Percentages of phenotypic classes are shown. **e-e’**. Mild hyperoxia alleviates cellular stress (TRE-NLS4xNG) caused by TOR overactivation (*UAS-Rheb*). Representative images are shown in **e** and quantification in **e’** (n ≥ 9 wing discs for each genotype, except for data of *hh-GAL4, tub-GAL80^TS^* > *UAS-Rheb* where n = 8). Depending on the normality of the data, a two-sided unpaired t-test or a two-sided unpaired Wilcoxon signed-rank test was used to assess statistical significance (* for p < 0.05, ** for p < 0.01, **** for p < 10^-4, n.s. for not significant).

## DISCUSSION

In this paper, we have shown that, even in normoxic conditions, developing imaginal discs of *Drosophila* become increasingly hypoxic as they grow towards their final size. A first hint came from transcriptomic analysis of precisely timed L3 discs, which suggest a progressive decrease in OXPHOS, with a commensurate increase in glycolysis. This observation motivated us to develop SyHREns, a transgenic reporter of HIF-1α activity, which can be used to estimate changes in oxygen level at cellular resolution. SyHREns responds rapidly (in less than 2.5 hours) to a mild decrease in environmental oxygen from normoxia (20.9%) to 17%, making it the most sensitive transgenic hypoxia indicator to date, while retaining an excellent dynamic range. Our evidence of gradually increasing hypoxia combined with volumetric measurements of precisely timed imaginal discs suggests a correlation between growth rate and oxygen availability. As the terminal electron acceptor of nutrient oxidation, oxygen is needed for the efficient production of ATP, which fuels sustained growth^82^. As such, oxygen is analogous to a resource (like nutrients) in the logistic model that describes the evolution of population size under limited resources^14,83^. Therefore, it is conceivable that limited oxygen availability could account for growth deceleration in wing imaginal discs and perhaps in a wide range of biological tissues. It will nevertheless be difficult to establish a causal relationship, at least until a method is developed to clamp intracellular oxygen at specific levels.

Intracellular oxygen levels are determined by supply and demand. Growth contributes to demand, albeit indirectly, via ATP consumption, and is therefore expected to decrease available oxygen, in accordance with the previously documented post-translational effect of insulin/TOR pathway on Sima/HIF-1α^60,84,85,86^. It was therefore predictable that increasing TOR signalling, e.g. by *Rheb* overexpression, would overwhelm oxygen supply and trigger detectable hypoxia. Nevertheless, to our knowledge, growth-induced hypoxia under physiological conditions has not been quantitatively documented in metazoans, perhaps because of the lack of a sensitive hypoxia sensor. The non-autonomous effect of overactive TOR signalling on systemic oxygen supply is also notable and may explain the earlier report that compartmental activation of TOR signalling leads to non-autonomous growth reduction in the genetically unaffected compartment^87^. It is clear therefore that excess growth triggers hypoxia. As our results show, <u>normal</u> growth during development also puts pressure on oxygen supply since growing imaginal discs display hallmarks of hypoxia, such as a metabolic shift away from OXPHOS^40,41^ and a dampening of TOR activity through transcriptional activation of *scyl/REDD1*^70,71,78^. We conclude that both physiological and pathological TOR signalling puts pressure on oxygen supply.

On the supply side, animals rely exclusively on environmental oxygen, which is normally relatively stable. Before being imported by tissues and cells, environmental oxygen is distributed throughout the body by the circulatory system in vertebrates and the tracheal system in insects. Evidence from *Manduca sexta*, a holometabolous insect like *Drosophila*, shows that the tracheal system only expands between larval instars^88^. Therefore, during each instar, organismal oxygen supply is unlikely to scale with individual growing tissues and organs. This could explain the observed rising imaginal disc hypoxia during L3, but only in part. Oxygen transport to tissue and cells must also be limiting since P compartment-specific Rheb-induced hypoxia does not readily equilibrate with that in the A compartment. In summary, steady-state intracellular oxygen is determined by growth-induced demand, supply from the environment, and the efficiency of the distribution system to individual cells. We envision that the distribution system will be subject, at least in part, to genetic control, enabling different tissues and organs to access oxygen according to their target growth rate and final size. However, all things being equal, it is expected that at constant environmental oxygen cellular oxygen tension will decline and hence growth will decelerate.

Our study highlights that growth exerts negative feedback on itself: TOR signalling activates Sima/HIF-1α, which in turn suppresses TOR signalling. Artificially boosting TOR signalling leads to cellular stress, as indicated by activation of JNK signalling. A likely scenario is that excessive oxygen demand leads to shortage, and hence depletion of terminal electron acceptors for OXPHOS and production of reactive oxygen species (ROS), which are known to damage DNA, proteins, and lipids, thus triggering cellular stress. As we have shown, such stress is exacerbated by *sima/HIF-1α* knockdown and alleviated by a mild increase in environmental oxygen, indicating that excess TOR signalling causes cellular stress by putting impossible demands on available oxygen supply. *Sima/HIF-1α* knockdown in otherwise wild type animals leads to mild cellular stress, suggesting that normal growth operates near the limit of available oxygen. Moreover, the wing phenotype of *sima/HIF-1α* mutants is partially reverted by inhibition of TOR activity (i.e. reducing demand). Collectively, these observations suggest that the Sima/HIF-1α-mediated feedback loop ensures that during normal development, growth-induced oxygen consumption is not allowed to exceed available supply.

## DATA AVAILABILITY

The RNA-Seq data will be publicly available at the Gene Expression Omnibus before publication. FlyBase ID and names for the genes identified by RNA-Seq are available at FlyBase (http://flybase.org/). All data supporting the findings of this study are available within the article or from the corresponding authors upon reasonable request.

## ACKNOWLEDGMENTS

We are grateful to Birgit Arne and Nic Tapon for allowing us to use their *ubi-NLS4xNG* strain. We also thank Ana Bolhaqueiro for characterisation of the *TRE-NLS4xNG* strain. The Advanced Light Microscopy Facility (CALM) at the Francis Crick Institute provided help with imaging and the Advanced Sequencing Facility (ASF) performed library preparation and RNA-Seq (Deb Jackson and Robert Goldstone). Ben Nicholls-Mindlin provided comments on the manuscript. YZ was the recipient of a PhD studentship from the Francis Crick Institute. The initial phase of this work was funded in part by an EMBO fellowship to GPM (ALTF 238-2018) and a Wellcome Trust Investigator Award to JPV (206341/Z/17/Z). The project was completed with core funding from the Francis Crick Institute to JPV (FC001204). The Francis Crick Institute receives its core funding from Cancer Research UK, the UK Medical Research Council, and the Wellcome Trust.

## ONLINE METHODS

### *Drosophila* strains and husbandry

*Drosophila* strains used in this study are summarised in Table 1. Flies were reared on fly food containing 5.5 g/L agar, 50 g/L glucose, 30 g/L wheat flour, 70 g/L yeast, 4.5 mL/L Propionic acid (Sigma-Aldrich, P5561), 19.5 mL/L Nipagin/Bavistin solution, 2.5 mL/L Penicillin/Streptomycin solution (Sigma-Aldrich, P4333) and ultrapure water. Unless otherwise noted, flies were maintained at 25°C in a Sanyo incubator with 12-hour light/dark cycles.

**Table 1.**
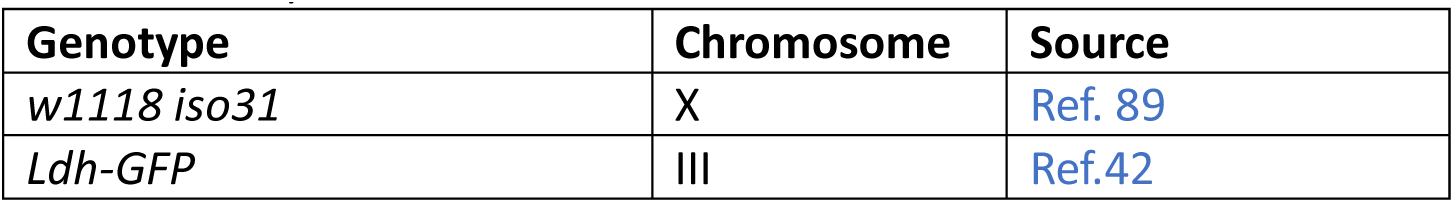
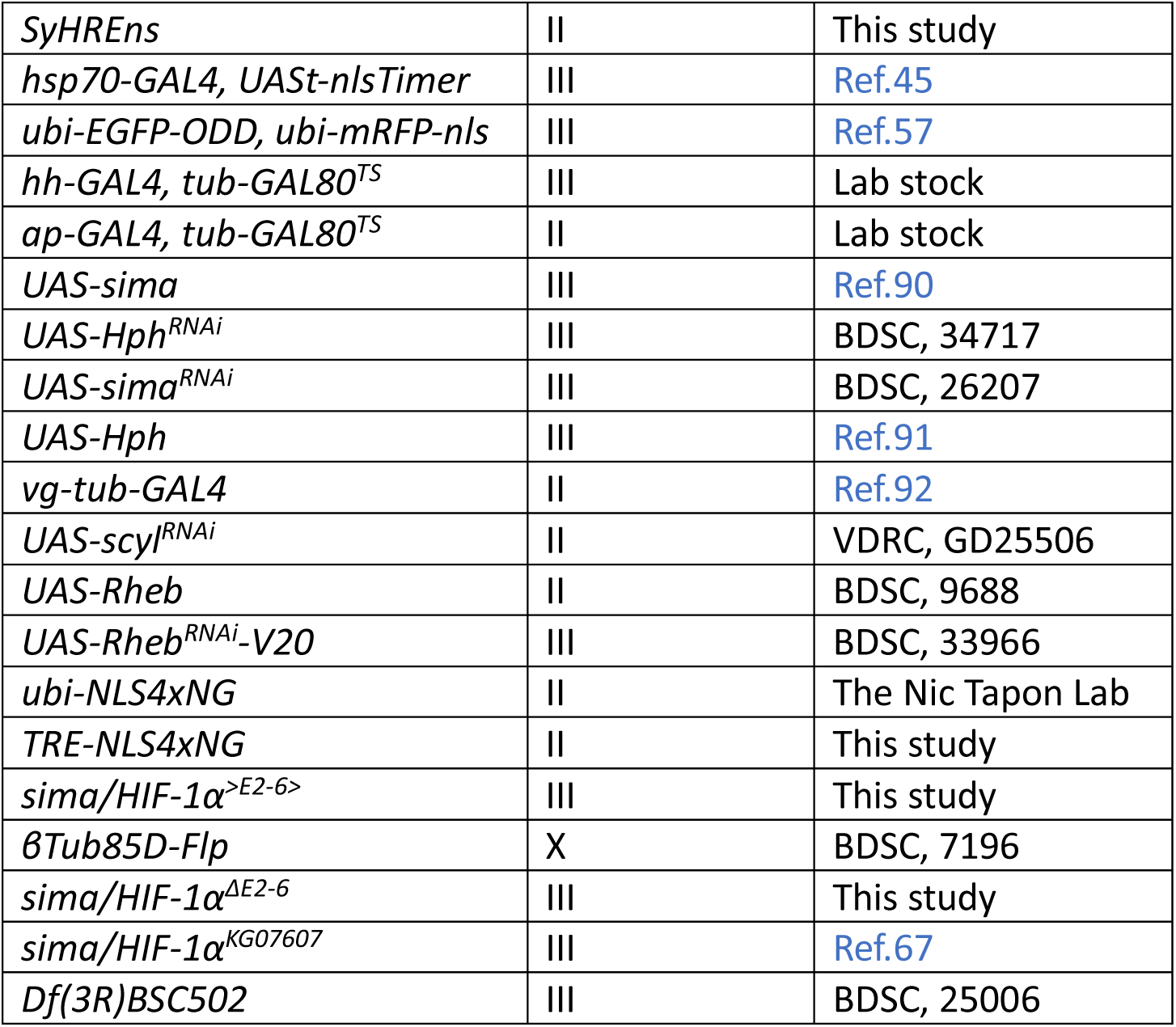
*Drosophila* strains.

### Generation of TRE-NLS4xNG

4x TPA response element (TRE)^93^ was inserted upstream of a minimal heat shock (hs) promoter that drives the expression of a NeonGreen tetramer with a nuclear localization signal (NLS4xNG)^20^. To enhance translation, a myosin heavy chain intervening sequence (IVS) and a synthetic AT-rich 21-bp sequence (Syn21) were placed upstream, and a highly-efficient p10 polyadenylation (polyA) signal was placed downstream of NLS4xNG. pJFRC81-10XUAS-IVS-Syn21-GFP-p10^66^ was a gift from Gerald Rubin (Addgene plasmid #36432). The plasmid injection to generate the TRE-NLS4xNG transgenic fly line was performed by The Fly Facility of the Francis Crick Institute.

### Generation of SyHREns (Synthetic HRE nuclear sensor)

5x hypoxia response element (HRE) was synthesised using the GeneArt Custom Gene Synthesis Services (Thermo Fisher Scientific). It consists of five 81-bp repeats. Each repeat contains a HRE2, a HRE3 and the intervening sequence, obtained from the 5’ upstream region of the sequence encoding *hph* isoform B that is upregulated in a Sima/HIF-1*α* dependent manner in response to hypoxia^65^. Subsequently, 4x TRE in the plasmid of TRE-NLS4xNG was replaced by 5x HRE. The plasmid was injected to BDSC, 9740 fly line by BestGene® to generate the SyHREns transgenic fly line.

### Generation of *sima/HIF-1α^ΔE2-6^* allele

A conditional allele *sima/HIF-1α^>E2-6>^* was initially created via CRISPR-Cas9 and homologous recombination-mediated repair as previously described^94^. Briefly, *sima/HIF-1α* was made conditional by substituting the region spanning exon 2 to exon 6 with an identical sequence flanked by two flippase recognition target (FRT) sites. CRISPR target sites were selected in unconserved regions, with one located in the intronic region between exon 1 and exon 2 (GGAATAGCTGCCGAGAACTTTGG) and another in the intronic region following exon 6 (CGTTAGTTGGTGGGTTGCAATGG). To generate the rescuing construct pTVmCherry-*sima/HIF-1α^>E2-6>^*, both the 5’ homology arm and the exon 2 to exon 6 of *sima/HIF-1α* flanked by two FRT sites (incorporated within the primers) were integrated upstream of a pax-mCherry selection marker through Gibson Assembly, whereas the 3’ homology arm was integrated downstream of the selection marker. The resulting pTVmCherry-*sima/HIF-1α^>E2-6>^* plasmid was co-injected with single guide RNAs (sgRNAs) into embryos from the nanos-Cas9 line. This injection was performed by the Fly Facility of the Francis Crick Institute. Successful candidates were first identified by pax-mCherry and subsequently confirmed by PCR. Following this, *β*-Tubulin at 85D (*β*Tub85D)-Flp (BDSC, 7196) was applied to excise the exon 2 to exon 6 in the male germline, thereby generating the *sima/HIF-1α^ΔE2-6^* allele.

### Quantification of wing disc volume

*Drosophila* larvae were selected at the L2-L3 transition (72 hours AEL) and subsequently allowed to develop for specific periods before being inverted in phosphate buffered saline (PBS), fixed in 4% Methanol-free Pierce^TM^ Formaldehyde (PFA) (Thermo Fisher Scientific, 28906) for 45 minutes (min) and stained with Invitrogen^TM^ Alexa Fluor^TM^ 647 Phalloidin (Thermo Fisher Scientific, A22287, concentration 1:267). To preserve the three-dimensional (3D) structure, the discs were dissected in PBS and mounted in 1% low melting point agar (Sigma-Aldrich, A9414) in PBS as previously described^26^. The mounted discs were immediately imaged with an upright Leica TCS SP5 confocal microscope using a 20x immersion objective with a z step of 1 μm. Since phalloidin labels all the actin-containing cells, it stains the entire wing disc tissue. The staining allows for the 3D reconstruction of the tissue using Imaris (RRID:SCR_007370) and relatively accurately measurement of the tissue volume.

### Quantification of wing disc growth rate

A logistic regression model was employed to describe the correlation between the volumes (*V*) of wing discs and their developmental stages over time (*t*). This model is particularly suited to scenarios where growth is initially rapid but decelerates as it approaches a maximum value, mirroring the physiological growth processes. To estimate the parameters for the model, a self-starting logistic regression model was applied. This approach automatically creates initial values for the parameters based on the provided data without manual initialisation, and it is described by the following equation:

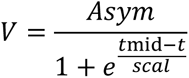

Where:

- *V* denotes the volume of the wing discs at developmental time *t*,
- *Asym* is the asymptotic maximum volume that the wing discs can achieve, analogous to the carrying capacity in traditional logistic models,
- *e* is the base of the natural logarithm,
- *t*_mid_ represents the time at which the volume is at half of its maximum value (*Asym/2*), serving as an inflection point in the context of growth,
- *scal* is a scaling parameter that affects the steepness of the curve, influencing how quickly the volume increases towards the asymptotic maximum.

The rate of change of volume with respect to time was determined by calculating *dV/dt*, the first derivative of *V*. This provides insight into the speed of volume increase at different developmental time points. The growth rate at a given time point was determined by dividing *dV/dt* by the instantaneous volume (*V_t_*), quantifying the relative increase in volume as a percentage of the instantaneous volume:

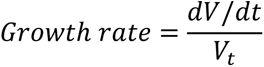

This measures the efficiency of volume increase relative to the size of the wing disc at different developmental time points, providing a detailed view of growth dynamics. All the analysis was conducted using RStudio^95^.

### Bulk RNA-Seq sample preparation

*Drosophila* larvae were selected at the L2-L3 transition (72 hours AEL) and subsequently allowed to develop for specific periods before the wing discs were dissected in PBS on ice within 20 min. Discs were then snap-frozen using dry ice and ethanol. The number of discs used for a single replicate at each time point are shown in Table 2. Three biological replicates were collected at each time point.

**Table 2.**
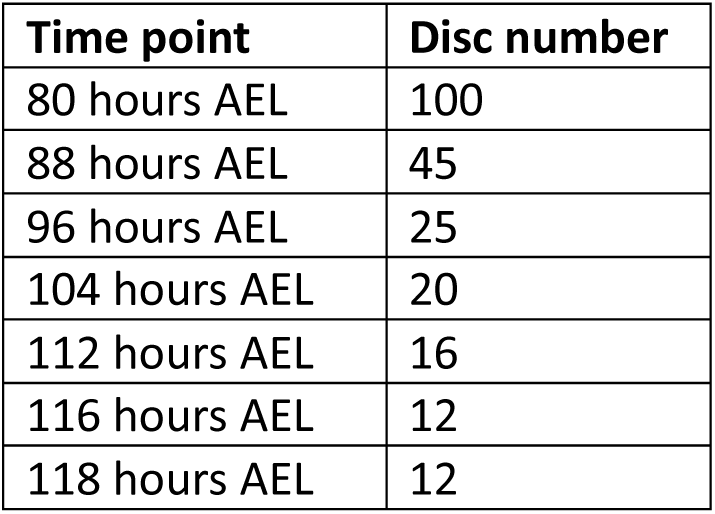
Number of wing discs used for RNA extraction per replicate.

| Time point | Disc number |
| --- | --- |
| 80 hours AEL | 100 |
| 88 hours AEL | 45 |
| 96 hours AEL | 25 |
| 104 hours AEL | 20 |
| 112 hours AEL | 16 |
| 116 hours AEL | 12 |
| 118 hours AEL | 12 |

After all the replicates were collected, RNA was extracted using the RNeasy Mini Kit (Qiagen, 74104), following the manufacturer’s protocol. Sequencing libraries were then prepared using the KAPA mRNA HyperPrep Kit (Roche, KK8581), following the manufacturer’s protocol. The Advanced Sequencing Facility of the Francis Crick Institute performed single-end 75 bp sequencing with 25 million reads per sample using a HiSeq 4000 System (Illumina).

### Bulk RNA-Seq analysis

The raw reads were analysed and normalised by the Bioinformatics and Biostatistics team of the Francis Crick Institute. The raw reads were quantified using version 3.0 of NFCore’s rnaseq pipeline^96^, with star-rsem as the alignment method against release 86 of Ensembl’s BDGP6 genome. Subsequently, these read counts were analysed using Bioconductor’s package DESeq2 version 1.30.1^97^ - differential genes were identified between time points by using a negative binomial model including main effects for time (as nominal variable) and experimental batch without any interaction, using Ward tests to assess the significance of shrunk^98^ fold-changes between pairs of time points, using a false-discovery threshold of 1%^99^.

Subsequently, to identify candidate genes potentially correlating with the decreasing growth rate, the Pearson correlation coefficient (*r*) between the changing transcript level over time and the decreasing growth rate was evaluated for each gene using RStudio^95^. Genes with *r* > 0.8 (increasing transcript level over time) or *r* < -0.8 (decreasing transcript level over time) were subjected to KEGG pathway analysis using the Database for Annotation, Visualisation and Integrated Discovery (DAVID)^100^. The identified KEGG pathway terms with a *p*-value < 0.05 were then subjected to graphic representation using RStudio^95^.

For heatmap representation, transcript level at each time point were further normalised to the maximum value observed over time for each gene. A distance matrix was generated using the Euclidean distance for continuous gene expression data. Subsequently, genes were clustered by assessing their similarities using the ward.D2 method^101^. All the analysis was conducted using RStudio^95^.

### Compartmental genetic perturbation

To achieve compartmental genetic perturbation in wing discs, either *hh-GAL4* or *apterous*(*ap*}*-GAL4* was used together with the temperature-sensitive *tubulin*(*tub)-GAL80^TS^*.

The larvae were reared at 18°C, the *GAL80^TS^* permissive temperature for five days AEL. To activate the compartmental genetic perturbation, the larvae were transferred to 29°C, the *GAL80^TS^* restrictive temperature. At this temperature, *GAL80^TS^* is inhibited, allowing the *GAL4*-driven gene expressions. Following two days at 29°C, the larvae were inverted and fixed for imaging.

### Validation of SyHREns by hypoxia treatment

SyHREns homozygous larvae were selected at the L2-L3 transition (72 hours AEL) and allowed to develop for an additional 24 hours to reach 96 hours AEL. Subsequently, the larvae were transferred into vials (15 larvae per vial) containing 1 mL 0.5% low melting point agarose (Sigma-Aldrich, A9414) in MilliQ deionised water. The vials were then evenly divided into two groups: the treatment group was placed in a hypoxia chamber set to a specific oxygen tension in a Sanyo MIR-154 incubator at 25°C; the control group was placed in a similar chamber within the same incubator but exposed to environmental normoxia. Both chambers were humidified using wet paper towels. After 2.5 hours of treatment, the larvae were inverted within 20 min and fixed for imaging. Oxygen tension in the hypoxia chamber was precisely maintained by an oxygen controller (Coy Laboratory Products) connected to nitrogen and oxygen gas cylinders. The oxygen chambers were custom-built by the Alex Gould Lab and the Making Lab of the Francis Crick Institute.

### Compartmental TOR overactivation under hypoxia or hyperoxia

For compartmental TOR overactivation under hypoxia or hyperoxia conditions, the larvae were initially reared at 18°C for five days AEL on fly food under environmental normoxia. The treatment group were then transferred to the hypoxia or hyperoxia chamber set to specific oxygen tension at 29°C. Meanwhile, the control group were transferred to a similar chamber at 29°C but kept under environmental normoxia. After two days of *GAL80^TS^* inactivation under either hypoxia or hyperoxia, the larvae were inverted and fixed for imaging.

### Larval oxygenation assay

Larvae expressing *hs-GAL4 > UASt-nlsTimer* were selected at L2-L3 transition (72 hours AEL). Both 72 and 84 hours AEL larvae were simultaneously subjected to a heat shock treatment at 37°C for 1 hour to induce a pulse of *nlsTimer* expression. Subsequently, nlsTimer was allowed to mature for 31 hours *in vivo* in both age groups of larvae. As described by Lehner and colleagues^45^, the larvae were mounted in 90% glycerol in PBS and immobilised at -20°C for 20 min before being imaged with an upright Leica TCS SP5 confocal microscope using a 10x immersion objective with a z step of 1 μm.

### Immuno- and endogenous fluorescence analysis

*Drosophila* larvae were inverted in PBS, fixed in 4% PFA for 30 min, washed in PBS for 10 min and then permeabilised in PBS with 0.1% Triton X-100 (0.1% PBST) for 30 min. For immunofluorescence imaging, the samples were blocked in 5% Normal Goat Serum (NGS) (Thermo Fisher Scientific, 01-6201) in 0.1% PBST for 30 min before being incubated in primary antibodies diluted in 0.1% PBST at 4°C overnight. Following three washes in 0.1% PBST for 15 min each, the samples were incubated in secondary antibodies diluted in 0.1% PBST at room temperature for 2 hours. After a further three 15-min washes in 0.1% PBST, the samples were incubated in Vectashield® with DAPI (2BScientific, H-1200-10) at 4°C overnight. Antibodies used in this study are shown in Table 3. For endogenous fluorescence imaging only, the samples were directly incubated in Vectashield® with DAPI at 4°C overnight post-permeabilisation. Subsequently, the wing discs or eye discs were dissected in PBS, mounted in Vectashield® with DAPI, and imaged with either an upright Leica TCS SP5 or an inverted Leica Falcon SP8 confocal microscope using a 40x immersion objective with a z step of 0.7 μm.

**Table 3.** Antibodies.

| Antibody | Host | Source | Concentration |
| --- | --- | --- | --- |
| pS6 | Rabbit | A gift from Jongkyeong Chung, Ref. 102 | 1:5000 |
| pS6 | Rabbit | A gift from Felipe Karam Teixeira, Ref. 103 | 1:500 |
| S6 (54D2) | Mouse | Cell Signaling Technology, 2317, RRID:AB_2238583 | 1:100 |
| Ci | Rat | DSHB, 2A1, RRID:AB_2109711 | 1:100 |
| β-Tubulin | Mouse | DSHB, E7, RRID:AB_528499 | 1:500 |
| Alexa Fluor™ Plus 488, anti-rabbit | Goat | Thermo Fisher Scientific, A32731, RRID:AB_2633280 | 1:1000 |
| Alexa Fluor™ Plus 555, anti-rabbit | Goat | Thermo Fisher Scientific, A32732, RRID:AB_2633281 | 1:1000 |
| Alexa Fluor™ Plus 488, anti-mouse | Goat | Thermo Fisher Scientific, A32723, RRID:AB_2633275 | 1:1000 |
| Alexa Fluor™ 647, anti-rat | Goat | Thermo Fisher Scientific, A-21247, RRID:AB_141778 | 1:1000 |

### Quantification of fluorescence intensity

For wing disc fluorescent signals, quantification of intensity was specifically focused on the wing pouch, distinguishable by the folds surrounding this region. The AP boundary within the wing pouch was identified by anti-Ci staining. Using Fiji, both the wing pouch and the AP boundary were outlined manually and a threshold excluding the background signals was applied to create a binary mask. The resulting regions of interest (ROIs) were used to determine the average fluorescence intensity either across the entire wing pouch or within each compartment. Unless otherwise noted, a single confocal plane that optimally captures the entire wing pouch was used for quantification and image representation.

For nuclear florescent signals, thresholding was applied to isolate the nuclei as a binary mask. The resulting ROIs were used to determine the average fluorescence intensity across the entire wing pouch, within each compartment, or throughout the larva.

### Quantification of TRE area

Using Fiji, both the wing pouch and the AP boundary were outlined manually, and the area of each compartment was determined. To segment the TRE positive nuclei, thresholding was applied to isolate the nuclei as a binary mask. The resulting ROIs were used to determine the total TRE area in each compartment. TRE area was calculated as a percentage of the compartment area occupied by the TRE positive nuclei respectively.

### Characterisation of SyHREns

The characterisation of SyHREns involved assessing its response to varying oxygen tensions. To determine the sensitivity and dynamic range of SyHREns in the wing pouches, 96 hours AEL larvae were exposed to external oxygen levels ranging from 5.0% to 20.9% for a duration of 2.5 hours.

### Mathematical modelling

For external oxygen levels between 5.0% and 15.0%, the average SyHREns intensities were normalised relative to their respective normoxia controls and plotted against the corresponding external oxygen levels (O_2_). To quantify the correlation between normalised SyHREns intensity (*I*) and O_2_ within this range, a linear regression model was applied, described by the following equation:

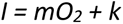

Where:

- *I* represents the normalised average SyRHEns intensity,
- *m* is the slope of the regression line, indicating the change in intensity with external oxygen levels (O_2_),
- *O_2_* is the external oxygen level,
- *k* is the y-intercept of the regression line.

In contrast, for external oxygen levels between 18.0% and 20.9%, the average SyHREns intensities (*I_c_*) were found to be independent of external oxygen levels (O_2_). This observation suggests SyHREns does not respond to O_2_ within this higher range. Consequently, a linear regression model with a slope of zero (*m* = 0) was applied, described by the following equation:

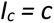

Where:

- *I_c_* represents the constant average SyRHEns intensity under 18.0% - 20.9% O_2_,
- *c* is the y-intercept of the regression line, denoting the constant value of average SyHREns intensity within a higher range of O_2_.

### Sensitivity of SyHREns

The sensitivity of SyHREns, defined as the mildest hypoxia to which SyHREns can respond within the 2.5-hour exposure, was determined by identifying the intercept of the two linear regression functions. This intercept occurs at O_2_ = 17.09%, marking the transition point below which SyRHEns begins to respond to decreasing oxygen levels.

### Dynamic range of SyHREns

The dynamic range of SyHREns, which quantifies the extent of its response from the sensitivity threshold down to 5.0% O_2_, was calculated as the fold-increase in intensity from 17.09% to 5.0% O_2_. This analysis revealed that SyHREns demonstrates an approximately 4- fold increase in intensity across its responsive oxygen range, indicating its capability to dynamically report changes in external oxygen level under specified experimental conditions and imaging with an upright Leica TCS SP5 confocal microscope.

### Western blot analysis of wing discs

Larvae were selected at the L2-L3 transition (72 hours AEL) and subsequently allowed to develop for specific periods before the wing discs were dissected in PBS on ice and lysed in 1x RIPA Lysis Buffer (Merck Millipore, 20-188) with 1x Ethylenediaminetetraacetic acid (EDTA)-Free Halt^TM^ Protease Inhibitor Cocktail (Thermo Fisher Scientific, 87785). The lysates were then subjected to 15 cycles of sonication (20 seconds on, 20 seconds off) at a low frequency, using the Bioruptor® Plus (Diagenode, B01020001) at 4°C. The total protein concentration in each lysate was determined using the Pierce^TM^ Bicinchoninic Acid (BCA) Protein Assay Kit (Thermo Fisher Scientific, 23227). Subsequently, the samples were denatured in a mixture of 1x Invitrogen^TM^ Bolt^TM^ LDS Sample Buffer (Thermo Fisher Scientific, B0007) and 0.25x Invitrogen^TM^ Bolt^TM^ Sample Reducing Agent (Thermo Fisher Scientific, B0009) at 95°C for 5 min and cooled to room temperature. They were then loaded (4 μg total protein per well) alongside the PageRuler^TM^ Plus Prestained NIR Protein Ladder (Thermo Fisher Scientific, 26619) onto the Invitrogen^TM^ 4-12% Bolt^TM^ Bis-Tris Plus Mini Protein Gels (Thermo Fisher Scientific, NW04120BOX, NW04122BOX or NW04125BOX) and subjected to electrophoresis using 1x Invitrogen^TM^ MES SDS Running Better (Thermo Fisher Scientific, B0002). Following the electrophoresis, proteins were transferred from the gel onto the Immobilon®-FL PVDF Membrane (Merck Millipore, IPFL00010) in wet conditions at 120 Volts for 70 min using an Invitrogen^TM^ Mini Gel Tank (Thermo Fisher Scientific, A25977). After that, the membranes were first blocked in 5% skimmed milk in PBS with 0.1% Tween- 20 (0.1% PBSTw) for 30 min before being incubated in primary antibodies diluted in 0.1% PBSTw with 5% skimmed milk at 4°C overnight. Following three washes in 0.1% PBSTw for 15 min each, the membranes were incubated in secondary antibodies diluted in 0.1% PBSTw with 5% skimmed milk and 0.01% sodium dodecyl sulfate (SDS) at room temperature for 2 hours. After a further three 15-min washes in 0.1% PBSTw with 0.01% SDS, the membranes were imaged using an Odyssey CLx Imaging System (LI-COR). The protein band intensities were quantified using Image Studio^TM^ (LI-COR). Antibodies used in this study are shown in Table 3.

### O-propargyl-puromycin (OPP) incorporation assay

The protein synthesis rate was determined using the Click-iT^TM^ Plus OPP Alexa Fluor^TM^ 594 Protein Synthesis Assay Kit (Thermo Fisher Scientific, C10457) as previously described^104^.

Briefly, larvae were selected at the L2-L3 transition (72 hours AEL) and subsequently allowed to develop for specific periods before being inverted in Gibco^TM^ Schneider’s *Drosophila* Medium (Thermo Fisher Scientific, 21720024) at room temperature within 15 min. The samples were then incubated with 1 μM OPP in Gibco^TM^ Schneider’s *Drosophila* Medium at room temperature for 15 min before being washed in PBS and fixed in 4% PFA. Following a wash in PBS, a 30-min permeabilisation in 0.1% PBST and a 30-min blocking in 5% NGS in 0.1% PBST, the samples were subsequently incubated with the Click-iT^TM^ reaction cocktail in the dark at room temperature for 30 min. After being washed once with the Click-iT^TM^ Reaction Rinse Buffer, the samples were incubated in Vectashield® with DAPI. The wing discs were then dissected in PBS, mounted in Vectashield® with DAPI, and imaged with an upright Leica TCS SP5 using a 40x immersion objective with a z step of 0.7 μm.

### Hybridization chain reaction (HCR) and image acquisition

The visualisation of mRNA in wing discs was performed using the HCR^TM^ IF Bundle (Molecular Instruments), following a protocol adapted from Patel and colleagues^105^. The initial preparation involved inverting larvae in PBS, fixing then in 4% PFA for 1 hour, followed by a 10-min wash in PBS and a 30-min permeabilisation in 0.1% PBSTw. Subsequently, the samples were pre-hybridised in 300 μL pre-warmed Probe Hybridisation Buffer at 37°C for 30 min, followed by an overnight incubation in 300 μL Probe Hybridisation Buffer containing 1 pmol of each probe set at 37°C. Both steps employed a thermoshaker set at 300 rpm. Post-hybridisation, the samples were subjected to four 15-min washes in 1 mL pre-warmed Probe Wash Buffer at 37°C, and then two 5-min washes in 600 μL 5x SSCT at room temperature. 100 μL Amplification Buffer containing 6 pmol of each hairpin was incubated at 95°C for 90 seconds before being cooled to room temperature in the dark. After the washes, the samples were pre-amplified in 400 μL pre-warmed Amplification Buffer at room temperature for 30 min, followed by an overnight incubation in 100 μL hairpin-containing Amplification Buffer at room temperature in the dark. Both steps employed a thermoshaker set at 300 rpm. Post-amplification, the samples were subjected to two 5-min and two 30-min washes in 400 μL 5x SSCT at room temperature, followed by three 10-min washes in PBS. After that, the samples were incubated in Vectashield® with DAPI at 4°C overnight before the wing discs were dissected in PBS, mounted in Vectashield® with DAPI, and imaged with an inverted Leica Falcon SP8 confocal microscope using a 40x immersion objective with a z step of 0.7 μm.

### Rapamycin treatment

Embryos were laid either on fly food containing 2 μM Rapamycin (Merck Supelco, 37094) dissolved in ethanol or on fly food containing an equivalent volume of ethanol alone (0 μM Rapamycin). Following hatching, the larvae were allowed to feed and develop into adult flies before the penetrance of wing crumpling was quantified.

### Sample preparation and image acquisition of adult wings

Adult flies were incubated in a solution comprising 70% glycerol (v/v) and 30% ethanol (v/v) at room temperature overnight before being washed in distilled water. Wings were dissected in isopropanol (Thermo Fisher Scientific), mounted in Euparal (Agar Scientific), and dried at room temperature for 24 hours. Images were acquired using a Zeiss Axioplan 2 microscope with a Leica MC190 HD camera, using a 2.5x objective for general wing morphology or a 10x objective for detailed views of the wing hairs.

### Quantification of adult wing area

For quantification of wing area, thresholding was applied to isolate the adult wing as a binary mask using Fiji. The hinge region attached to the wing was manually removed from the binary mask to isolate the wing blade, allowing for measurement of wing area.

### Quantification of adult wing hair density

For assessment of wing hair density, a systematic approach was applied where four equally sized ROIs were selected within the wing, specifically between the longitudinal veins L2 and L3, L3 and L4, L4 and L5, and L5 and L6 using Fiji. In these ROIs, a threshold exclusive for the wing hair sockets excluding the background signals was applied to create a binary mask. The density of wing hairs was calculated based on the average count of hair sockets across the four ROIs for each wing, presented as the number of wing hairs per unit area.

### Timing of larval development

Embryos were laid on agar plates supplemented with yeast paste over 4-hour intervals from 8:00 to 20:00. L1 larvae were collected at 24 hours AEL from the agar plates and transferred into vials containing fly food. Each vial housed 35 larvae, with either six or nine vials prepared per genotype depending on the vitality of the fly lines (n = 210 or n = 315). Three days after the transfer, the number of larvae that had pupariated was recorded every 4-hour from 8:00 to 20:00 for three to five days. The proportion of larvae that transitioned to pupae within each population was then plotted against developmental time, generating a curve of larval developmental timing. The average timing of larval-pupal transition is defined by the time point where 50% of the population becomes pupae^106^.

## AUTHORS CONTRIBUTIONS

This project was conceived by GPM, JPV and YZ. CA built the TRE-NLS4xNG and the *sima/HIF-1α^>E2-6>^* strain. GK performed differential gene expression analysis from the raw RNA-Seq reads. YZ executed most of the experiments and data analysis, including bioinformatics. Data interpretation was done collectively by all authors. YZ wrote the first draft, which was subsequently modified and edited by all the authors.

---

**Extended Data Fig. 1:**
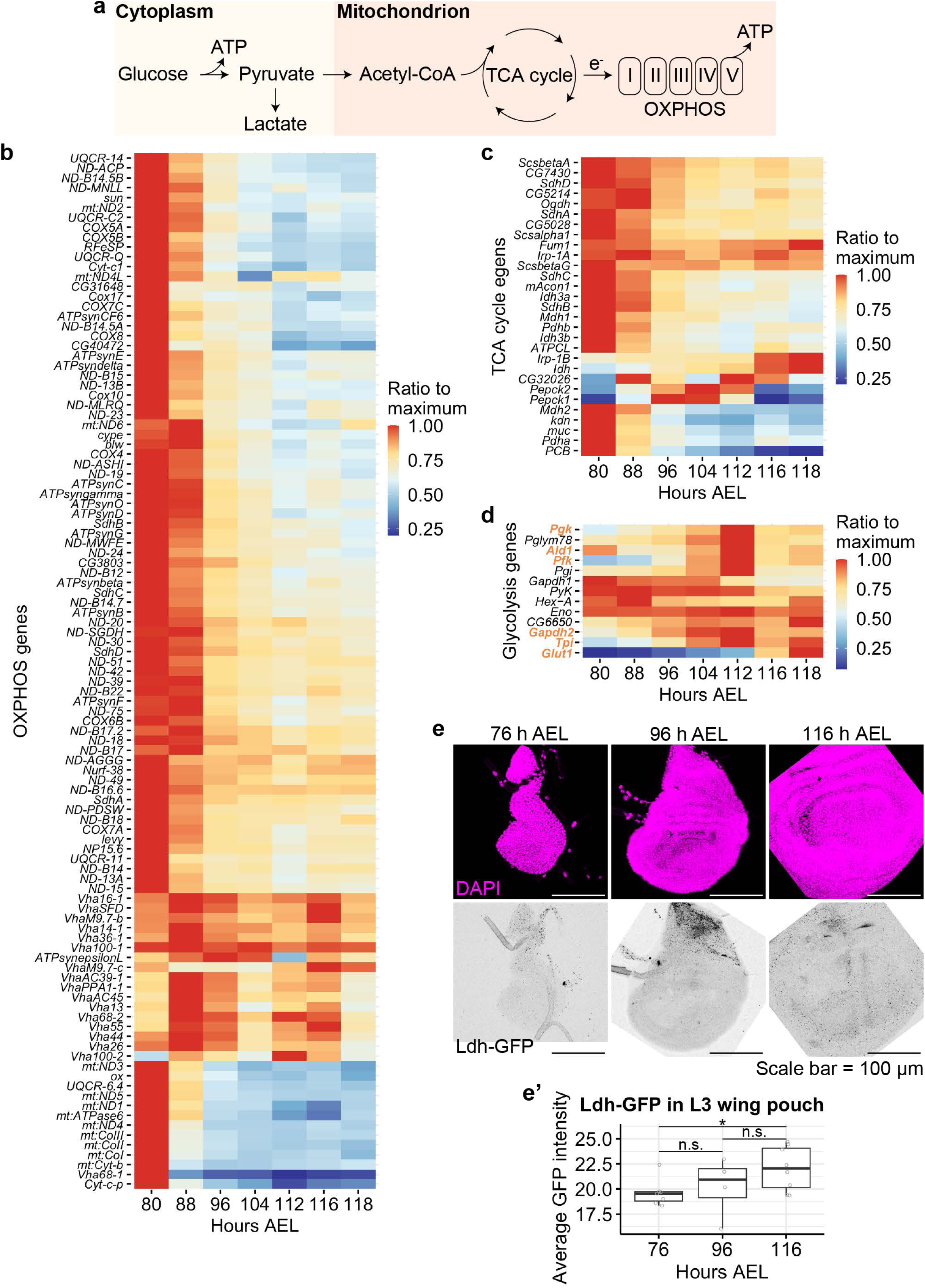
Transcriptomic analysis of L3 wing discs reveals transcriptional alterations typically associated with hypoxia. **a.** Schematic representation of aerobic metabolism with lighter shading highlighting anaerobic metabolism. **b.** Detailed transcriptional profile of the OXPHOS genes shown in Fig. 1f (with gene names). **c.** Transcriptional profile of TCA cycle genes expressed during L3. **d.** Transcriptional profile of glycolysis genes expressed during L3. Expression of *Pgk*, *Ald1*, *Pfk* and *Glut1* is known to be upregulated under hypoxia. **e-e’**. Elevation in *Ldh* expression in wing discs during L3, detected by Ldh-GFP. Representative images are shown in **e** and quantification in **e’** (for each time point, n ≥ 7 wing discs, except for data at 96 hours AEL where n = 4). After ensuring the normality of the data by performing a Shapiro-Wilk test, a two-sided unpaired t-test was used for statistical significance (* for p < 0.05, n.s. for not significant). For all the box plots in this and subsequent figures, the centre line denotes the median (50th percentile); the lower quartile (Q1) denotes the 25th percentile; the upper quartile (Q3) denotes the 75th percentile; the interquartile range (IQR) is defined as *Q*3 − *Q*1; the outliers are defined as data points that lie below *Q*1 − 1.5 × *IQR* or above *Q*3 + 1.5 × *IQR*.

**Extended Data Fig. 2:**
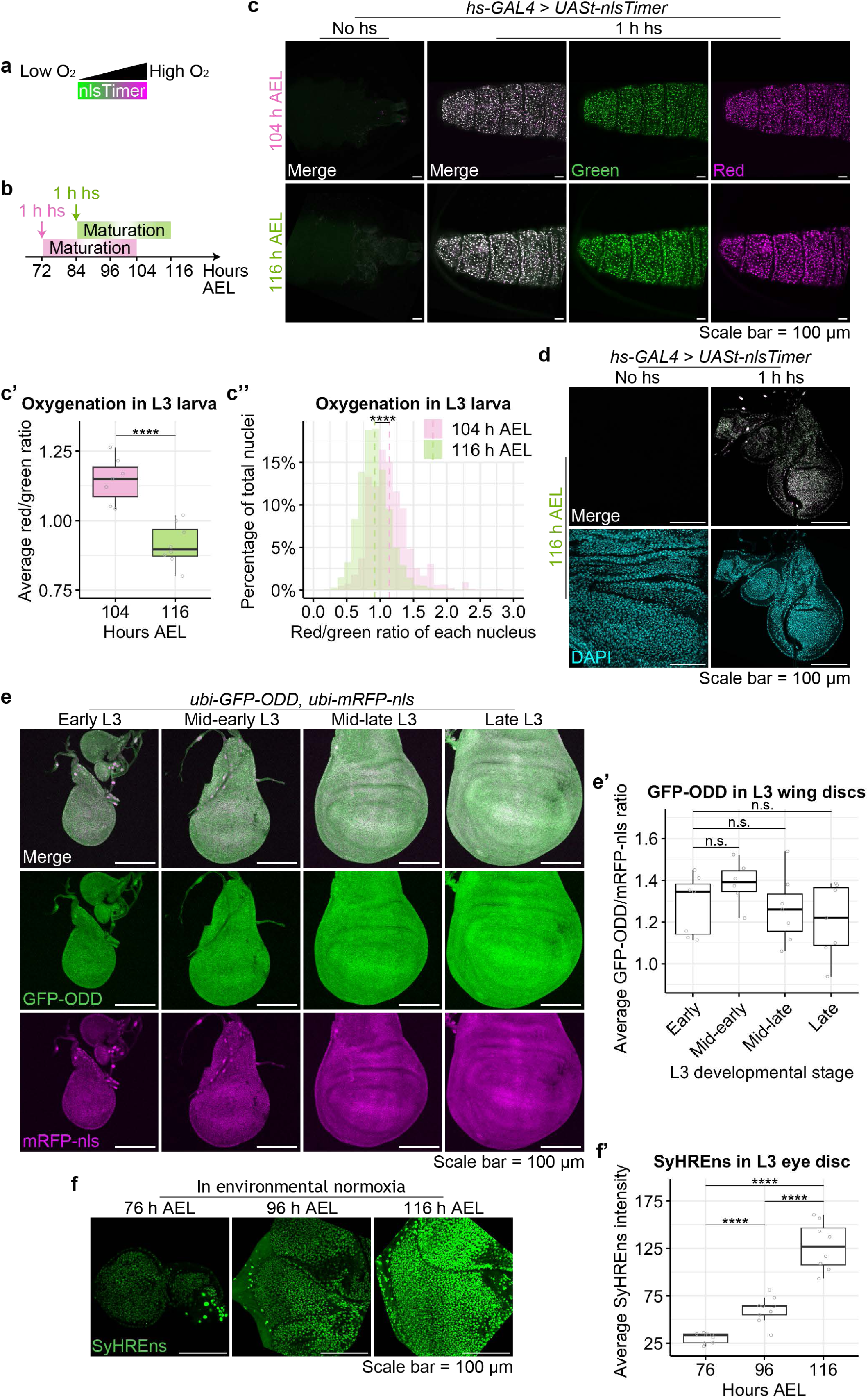
Transcriptomic analysis of L3 wing discs reveals changes normally associated with hypoxia. **a.** Schematic representation of nlsTimer’s maturation at different oxygen levels. **b.** Experimental protocol to compare oxygen levels in young and old L3 larvae with nlsTimer. **c-c’’**. Red and green signals from nlsTimer at 104 and 116 hours AEL. Representative images are shown in **c**. Quantification (**c’** and **c’’**) shows that the red signal is relatively higher than the green one in the younger discs (n ≥ 7 larvae for each time point). After ensuring the normality of the data with a Shapiro-Wilk test, a two-sided unpaired t-test was used to assess statistical significance in **c’** (**** for p < 10^-4). A two-sided unpaired Kolmogorov Smirnov test (K-S test) was performed to determine that the two histograms in **c’’** are statistically distinct (Distance = 0.3314, **** for p < 10^-4). **d**. Heat-shock-induced nlsTimer expression (with *hs-GAL4 > UASt-nlsTimer*) impairs growth in wing imaginal discs. **e-e’**. Fluorescence from GFP-ODD remains constant during L3. Representative images are shown in **e** and quantification in **e’** (n = 7 wing discs for each developmental stage, except for data at mid-early stage where n = 6). After ensuring the normality of the data, a two-sided unpaired t-test was used for statistical significance (n.s. for not significant). **f-f’**. SyHREns activity progressively rises in L3 <u>eye</u> discs under environmental normoxia. Representative images are shown in **f** and quantification in **f’** (n = 9 eye discs for each time point, except for data at 116 hours AEL where n = 8). After ensuring the normality of the data, a two-sided unpaired t-test was performed to assess statistical significance (**** for p < 10^-4).

**Extended Data Fig. 3:**
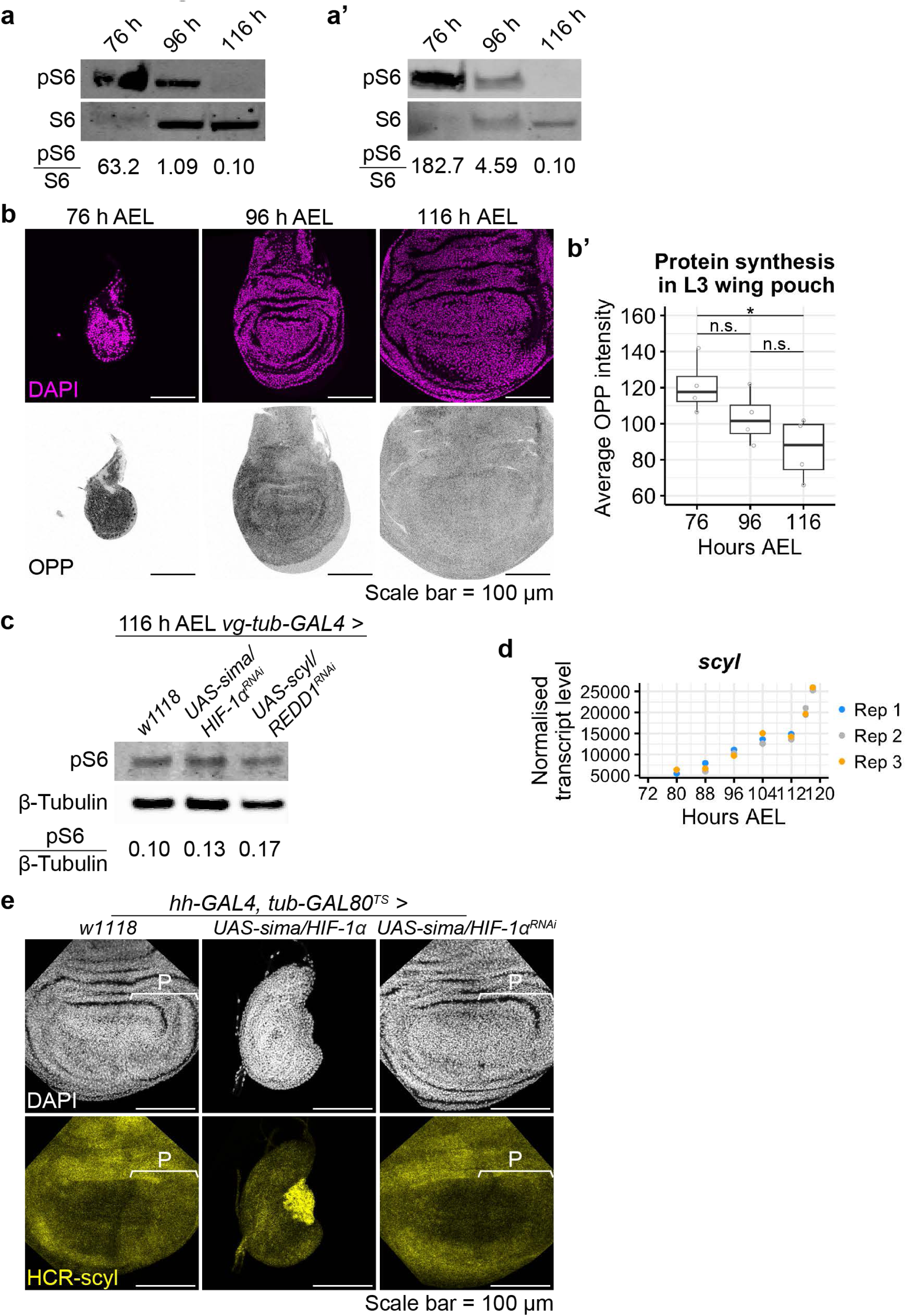
A regulatory link between Sima/HIF-1α, Scyl/REDD1 and TOR signalling. **a-a’**. Progressive decrease in TOR activity (as detected with anti-pS6) in developing wing discs during L3. pS6 was normalised as a ratio to total S6. Two independent experiments are show. **b-b’**. The rate of protein synthesis (detected by OPP incorporation) diminishes with age/size in L3 discs. Representative images are shown in **b** and quantification in **b’** (n = 4 wing discs for each time point). After ensuring the normality of the data with a Shapiro-Wilk test, a two-sided unpaired t-test was used to assess statistical significance (*for p < 0.05, n.s. for not significant). **c.** Western blot analysis shows a mild increase of pS6 levels upon whole disc knockdown of *sima/HIF-1α* or *scyl/REDD1* (*vg-tub-GAL4 > sima/HIF-1α^RNAi^* or *scyl/REDD1^RNAi^*). The pS6 signal, normalised with anti-β-Tubulin is indicated as a ratio under each lane. This is a repeat of the experiment shown in Fig. 3c. **d.** Bulk RNA-Seq of wing imaginal discs reveals rising level of *scyl/REDD1* mRNA during L3. **e.** Perturbations of *sima/HIF-1α* activity (*hh-GAL4 > sima/HIF-1α* or *sima/HIF-1α^RNAi^*) affect *scyl/REDD1* transcript levels, as assessed by hybridization chain reaction (HCR). The P compartment, where *hh-GAL4* is expressed, is marked by a bracket.

**Extended Data Fig. 4:**
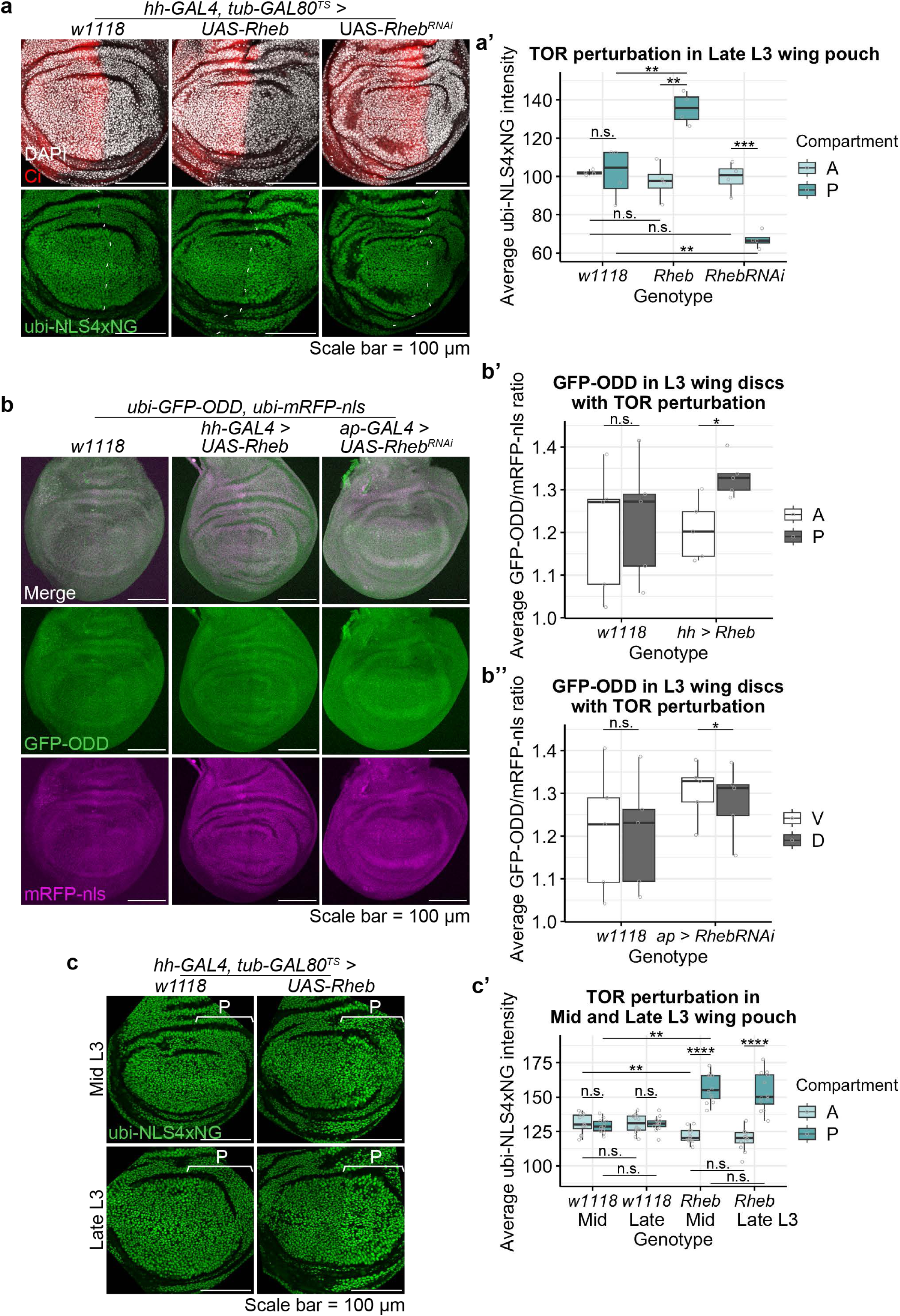
Effect of excess TOR activity on expression of a constitutive reporter gene and the GFP-ODD hypoxia reporter. **a-a’.** Perturbation of TOR activity autonomously affects the expression of a constitutive reporter gene (*ubi-NLS4xNG*) without detectable non-autonomous effect. Representative images are shown in **a** and quantification in **a’** (n = 4 wing discs for each genotype). After testing for normality with a Shapiro-Wilk test, a two-sided unpaired t-test was used to assess statistical significance (** for p < 0.01, *** for p < 0.001, n.s. for not significant). **b-b’’.** Perturbation of TOR activity affects the expression of the GFP-ODD hypoxia reporter autonomously. Representative images are shown in **b** and quantification in **b’** and **b’’** (n = 5 wing discs for each genotype). After ensuring the normality of the data, a two-sided unpaired t-test was used to assess statistical significance (* for p < 0.05, n.s. for not significant). **c-c’’**. Rheb overexpression similarly boosts the expression of a constitutive reporter gene (*ubi-NLS4xNG*). Representative images are shown in **c** and quantification in **c’** and **c’’**. (n = 6 wing discs for each condition). After ensuring the normality of the data, a two-sided unpaired t-test was used to assess statistical significance (** for p < 0.01, **** for p < 10^-4, n.s. for not significant).

**Extended Data Fig. 5:**
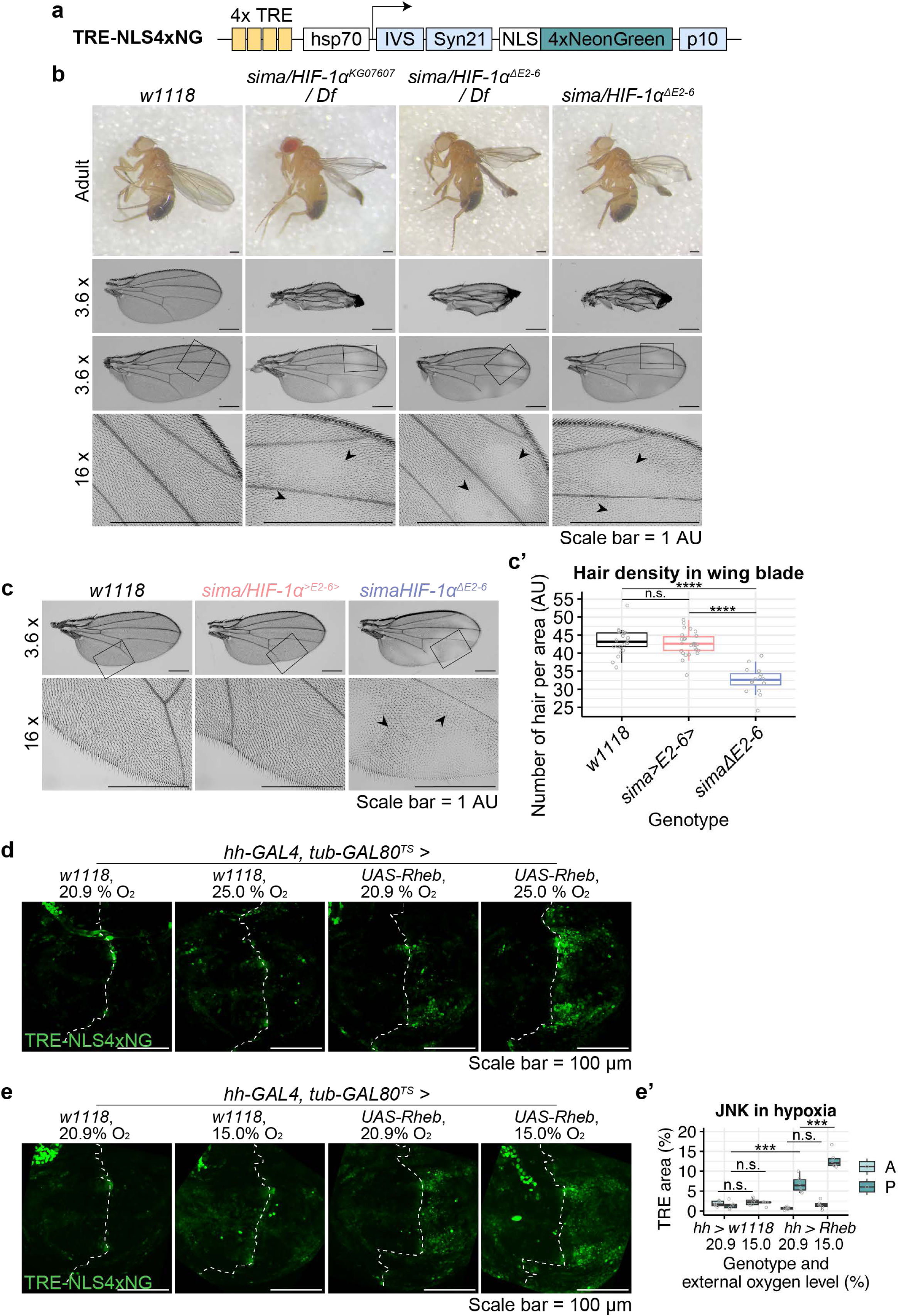
Effect of Sima/HIF-1α mutation or oxygen deprivation on morphological defect and cellular stress. **a.** Schematic representation of our JNK reporter, TRE-NLS4xNG, adapted from 12-O-Tetradecanoylphorbol-13-acetate (TPA) response element (TRE) reporters designed by Chatterjee and Bohmann (Chatterjee and Bohmann, 2012). TRE represents the consensus DNA binding of activator protein 1 (AP-1). **b.** Phenotypes of *Sima/HIF-1α* mutant wings. These phenotypes can be classified as mild (loss of hair in patches, black arrowheads + altered hair morphology and density, as shown in panel c below) or severe (‘crumpled’ wings, in addition to the hair phenotypes) Representative images are shown. For *sima/HIF-1α^ΔE2-6^* homozygotes, the frequency was 41.4% mild and 58.6% severe (see Fig. 5d). Neither phenotype was seen in wild type wings. Black rectangles in the 3.6 x images represent the areas of interest for 16 x magnification. **c-c’’**. Wing hair phenotypes of *Sima/HIF-1α* mutant flies. Representative images are shown in **c** and quantification in **c’** (n ≥ 24 adult wings for each genotype, except for data of *sima/HIF-1α^ΔE2-6^* homozygotes where n = 16). Crumpled wings are not shown since difficult to image at high magnification. Black rectangles in 3.6 x images represent the areas of interest for 16 x magnification. Black arrows point towards abnormally long and soft wing hairs. After ensuring the normality of the data with a Shapiro-Wilk test, a two-sided unpaired t-test was used to assess statistical significance (**** for p < 10^-4, n.s. for not significant). **d**. TOR-induced cellular stress (TRE-NLS4xNG) is exacerbated by 25% environmental oxygen tension. Representative images are shown in **e** (n ≥ 8 wing discs for each genotype). **e-e’**. TOR-induced cellular stress (TRE-NLS4xNG) is exacerbated by mild hypoxia. Representative images are shown in **e** and quantification in **e’** (n ≥ 5 wing discs for each genotype). Depending on the normality of the data, a two-sided unpaired t-test or a two-sided unpaired Wilcoxon signed-rank test was used to assess statistical significance (*** for p < 0.001, n.s. for not significant).

